# Tau polarizes an aging transcriptional signature to excitatory neurons and glia

**DOI:** 10.1101/2022.11.14.516410

**Authors:** Timothy Wu, Jennifer M. Deger, Hui Ye, Caiwei Guo, Justin Dhindsa, Brandon T. Pekarek, Rami Al-Ouran, Zhandong Liu, Ismael Al-Ramahi, Juan Botas, Joshua M. Shulman

## Abstract

Aging is a major risk factor for Alzheimer’s disease (AD), and cell-type vulnerability underlies its characteristic clinical manifestations. We have performed longitudinal, single-cell RNA-sequencing in *Drosophila* with pan-neuronal expression of human tau, which forms AD neurofibrillary tangle pathology. Whereas tau- and aging-induced gene expression strongly overlap (93%), they differ in the affected cell types. In contrast to the broad impact of aging, tau-triggered changes are strongly polarized to excitatory neurons and glia. Further, tau can either activate or suppress innate immune gene expression signatures in a cell type-specific manner. Integration of cellular abundance and gene expression pinpoints Nuclear Factor Kappa B signaling as a potential marker for neuronal vulnerability. We also highlight the conservation of cell type-specific transcriptional patterns between *Drosophila* and human postmortem brain tissue. Overall, our results create a resource for dissection of dynamic, age-dependent gene expression changes at cellular resolution in a genetically tractable model of tauopathy.

## Introduction

Alzheimer’s disease (AD) is a progressive neurodegenerative disorder characterized by extracellular amyloid-beta neuritic plaques and intracellular tau neurofibrillary tangles (DeTure and Dickson, 2019; Scheltens et al., 2021). Tau neuropathologic burden is strongly correlated with cognitive decline, synaptic loss, and neuronal death (Arriagada et al., 1992; Braak and Braak, 1991; Gomez-Isla et al., 1997). Cell type-specific vulnerability is also an important driver of AD clinical manifestations, including its characteristic amnestic syndrome. Neurofibrillary tangles first appear in the transentorhinal cortex, entorhinal cortex, and CA1 region of the hippocampus, affecting resident pyramidal cells and excitatory glutamatergic neurons; cholinergic neurons of the basal forebrain are also particularly vulnerable (Mrdjen et al., 2019; Fu et al., 2018). Single-cell RNA-sequencing (scRNAseq) or single-nucleus RNA-sequencing (snRNAseq) are promising approaches to pinpoint cell type-specific mechanisms in AD, including those that may underlie neuronal vulnerability (Mathys, et al., 2019; Grubman et al., 2019; Lau et al., 2020; Zhou et al., 2020). Emerging data highlight altered transcriptional states and/or cell proportions for vulnerable versus resilient neurons, including excitatory or inhibitory neurons, respectively (Leng et al., 2021). snRNAseq profiles also implicate important roles for non-neuronal cells, including oligodendrocytes, astrocytes, and microglia (Grubman et al., 2019; Lau et al., 2020; Zhou et al., 2020). Microglial expression signatures, including genes with roles in innate immunity, are sharply increased in brains with AD pathology, and an important causal role in AD risk and pathogenesis is reinforced by findings from human genetics (Bohlen et al., 2019; Deczkowska et al., 2018; Bellenguez et al., 2022).

One important limitation to gene expression studies from human postmortem tissue is that only crosssectional analysis is possible, making it difficult to reconstruct dynamic changes over the full time-course of disease. In fact, age is the most important risk factor for AD, which develops over decades (Masters et al., 2015; Villemagne et al., 2013). Another potential challenge is identifying molecularly specific changes, since tau tangle pathology usually co-occurs with amyloid-beta plaques, along with other brain pathologies which can also cause dementia (e.g., lewy bodies or infarcts) (Kapasi et al., 2017). By contrast, animal models permit experimentally-controlled manipulations isolating specific triggers and their impact over time. For example, in mouse models of amyloid-beta pathology, scRNAseq and snRNAseq have implicated subpopulations of disease-associated microglia and astrocytes, and similar changes may also characterize brain aging (Keren-Shaul et al., 2017; Habib et al., 2020). Further, in tau transgenic models, activation of immune signaling by the nuclear factor kappa-light-chain-enhancer of activated B cells (NF-κB) transcription factor within microglia was found to be an important driver of pathologic progression (Wang et al., 2022). We recently characterized tau- and aging-induced gene expression changes in a *Drosophila melanogaster* tauopathy model, revealing perturbations in many conserved pathways such as innate immune signaling (Mangleburg et al., 2020). Over 70% of tau-induced gene expression changes in flies were also observed in normal aging. In this study, we deploy scRNAseq in *Drosophila* to map the cell-specific contributions of age- and tau-driven brain gene expression and identify NFκB signaling as a promising marker of neuronal vulnerability.

## Results

### Single cell transcriptome profiles of the tau transgenic *Drosophila* brain

Pan-neuronal expression of the human *microtubule associated protein tau* (*MAPT*) gene in *Drosophila* recapitulates key features of AD including misfolded and hyperphosphorylated tau, age-dependent synaptic and neuron loss, and reduced survival (Wittmann et al., 2001). We performed scRNAseq of adult fly brains in *tau^R406W^* transgenic *Drosophila* (*elav>tau^R406W^*) and controls (*elav-GAL4*), including animals aged 1, 10 or 20 days (Figures S1A and S1B). The GAL4-UAS expression system is used to express human tau in neurons throughout the central nervous system (CNS) (Brand and Perrimon, 1993). The R406W variant in *MAPT* causes frontotemporal dementia with parkinsonism-17, an autosomal dominant, neurodegenerative disorder with tau pathology (i.e., tauopathy). In flies, *tau^R406W^* causes similar neurodegenerative phenotypes to wild type *tau* but induces a more robust transcriptional response (Wittmann et al., 2001; Mangleburg et al., 2020). Following stringent quality control, transcriptome data from 48,111 single cells were available for our initial analyses, including from 6 total conditions (2 genotypes x 3 ages) (Figures S1C-S1E). In the integrated dataset, we identified 96 distinct cell clusters grouped by transcriptional signatures, and annotated cell-type identities to 59 clusters using available *Drosophila* brain scRNAseq reference data and established cell markers (Figures 1A and S2; Table S1). As expected, most cells in the fly brain were neurons (*CadN* expression, n=42,587), whereas glia were comparatively sparse (*repo* expression, n=5,524). Our dataset comprises a diverse range of cell types. Among all cell clusters, 49% were cholinergic neurons (*VAChT*), 20% were glutamatergic neurons (*VGlut*), 11% were GABAergic neurons (*Gad1*), and 7% were glia (*repo, Gs2*) (Figures 1B and S3). We also identified several major glial subtypes in the fly brain (Kremer et al., 2017), including astrocyte-like, cortex, chiasm, subperineurial, perineurial, and ensheathing glia, along with a group of circulating macrophages (hemocytes). Overall, our findings are consistent with results from prior scRNAseq studies of whole adult *Drosophila* brains (Davie et al., 2018).

**Figure 1.**
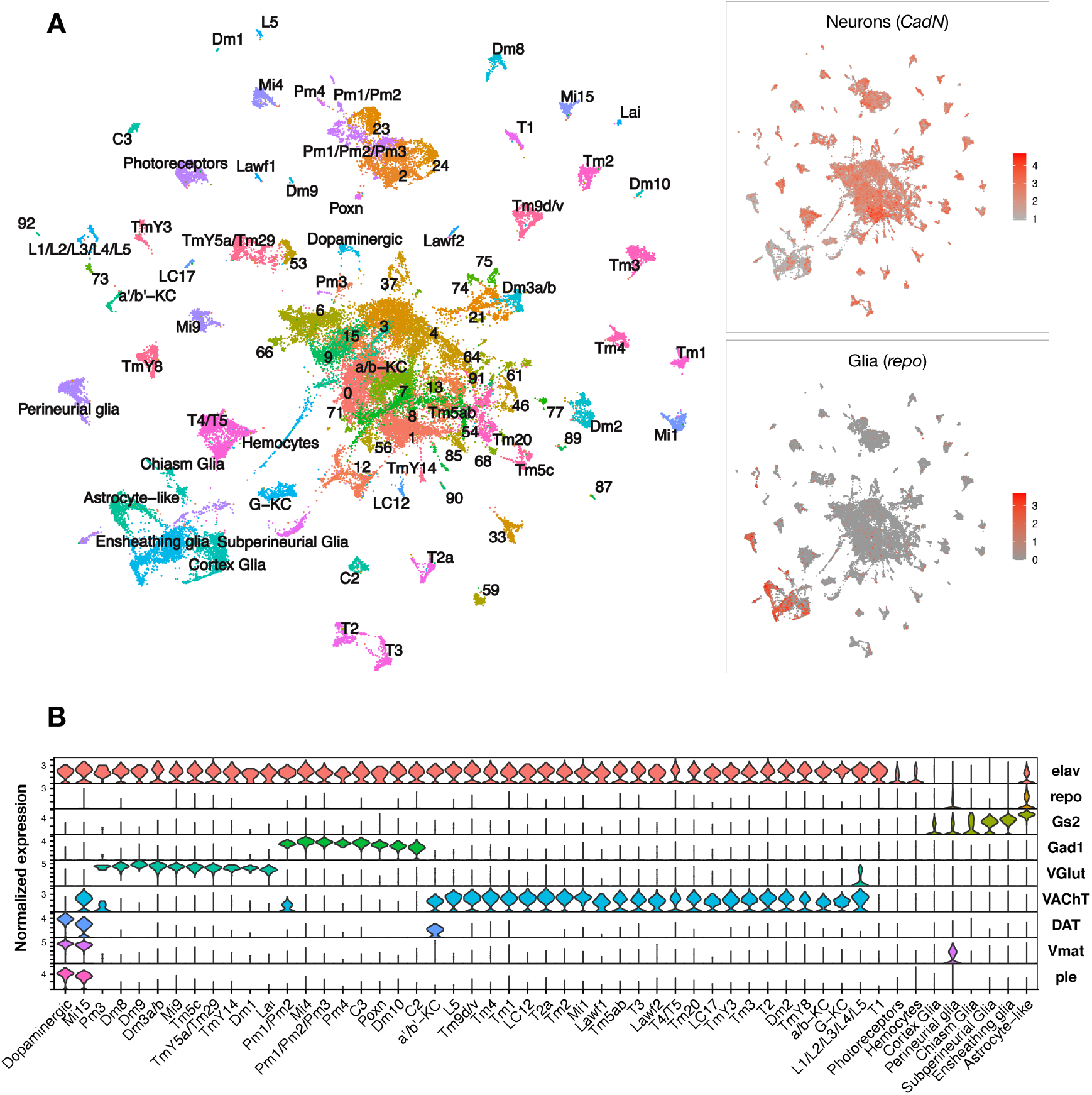
Single cell RNA-sequencing of the adult *Drosophila* brain. (A) Uniform manifold approximation and projection (UMAP) plot displays unsupervised clustering of 48,111 cells, including from control (*elav-GAL4 / +*) and *elav>tau^R406W^* transgenic animals (*elav-GAL4 ≥ +; UAS-tau^R406W^* / +) at 1-, 10-, and 20-days. Expression of neuron- and glia-specific marker genes, *CadN* and *repo*, respectively, are also shown. Cell cluster annotations identify heterogeneous optic lobe neuron types, including from the lamina (L1-5, T1, C2/3, Lawf, Lai), medulla (Tm/TmY, Mi, Dm, Pm, T2/3), and lobula (T4/T5, LC). Other identified neuron types include photoreceptors (*ninaC, eya*), dopaminergic neurons (*DAT, Vmat, ple*), and central brain mushroom body Kenyon cells (*ey, Imp, sNFP, trio*). (B) Violin plot showing cell type marker expression across annotated cell clusters. Selected markers include *Elav* (neurons), *repo/Gs2* (glia), *Gad1* (GABA), *VGlut* (glutamate), *VAChT* (acetylcholine), *DAT/Vmat/ple* (dopamine). See also Figures S1-S3 and Tables S1, S3 and S12-S13.

### tau drives changes in cell proportions in the brain

Leveraging our scRNAseq data and pooling longitudinal samples to permit robust comparisons, we first assessed how tau affects the relative abundance of cell type subpopulations in the adult brain. We found 16 neuronal and 6 glial clusters with statistically significant changes in cell abundance when comparing tau and controls (Figures 2A and 2B; Table S2). Cholinergic mushroom body Kenyon cell neurons in the central complex, which are important in learning and memory, were sharply reduced, consistent with prior studies of *Drosophila* tauopathy models (Mershin et al., 2004; Kosmidis et al., 2010). In fact, 7 excitatory neuronal clusters, including several cholinergic and glutamatergic cell types, demonstrated significant declines, whereas inhibitory neuronal subpopulations (e.g., Pm and Mi4 GABAergic cells in the visual system) appeared resilient to tau. Conversely, cluster 12 cells appeared more abundant in tau flies; this non-annotated cell type was enriched for neuroendocrine expression markers, *Ms* and *Hug*, as well as a regulator of synaptic plasticity, *Arc1* (Table S3). Interestingly, several glial cell types also appeared increased in the brains of tau animals. Ensheathing glia, which showed the largest potential increase, are localized to neuropil in the fly brain and mediate phagocytosis following neuronal injury (Doherty et al., 2009; Freeman, 2015). In order to confirm these observations, which were based on pooled data across timepoints, we generated additional scRNAseq profiles from 10-day old *elav>tau^R406W^* and control flies in triplicate samples (69,128 cells; Figure S4A). Overall, 13 out of the 22 significant cell abundance changes were also observed in this replication dataset, including the sharp reduction of excitatory neurons (e.g., Kenyon cells), and the increase in multiple glial clusters (e.g., ensheathing glia) (Figure S4B; Table S2). Non-replicated changes in cell type abundance may be driven by data from earlier (1-day) or later (20-day) timepoints (Figure 2B). Many cell type-proportion changes were also recapitulated based on computational deconvolution of available bulk-tissue RNAseq from *tau^R406W^* and control flies at 1-, 10-, and 20-days by using an independent, published scRNAseq reference dataset (Figure S5).

**Figure 2.**
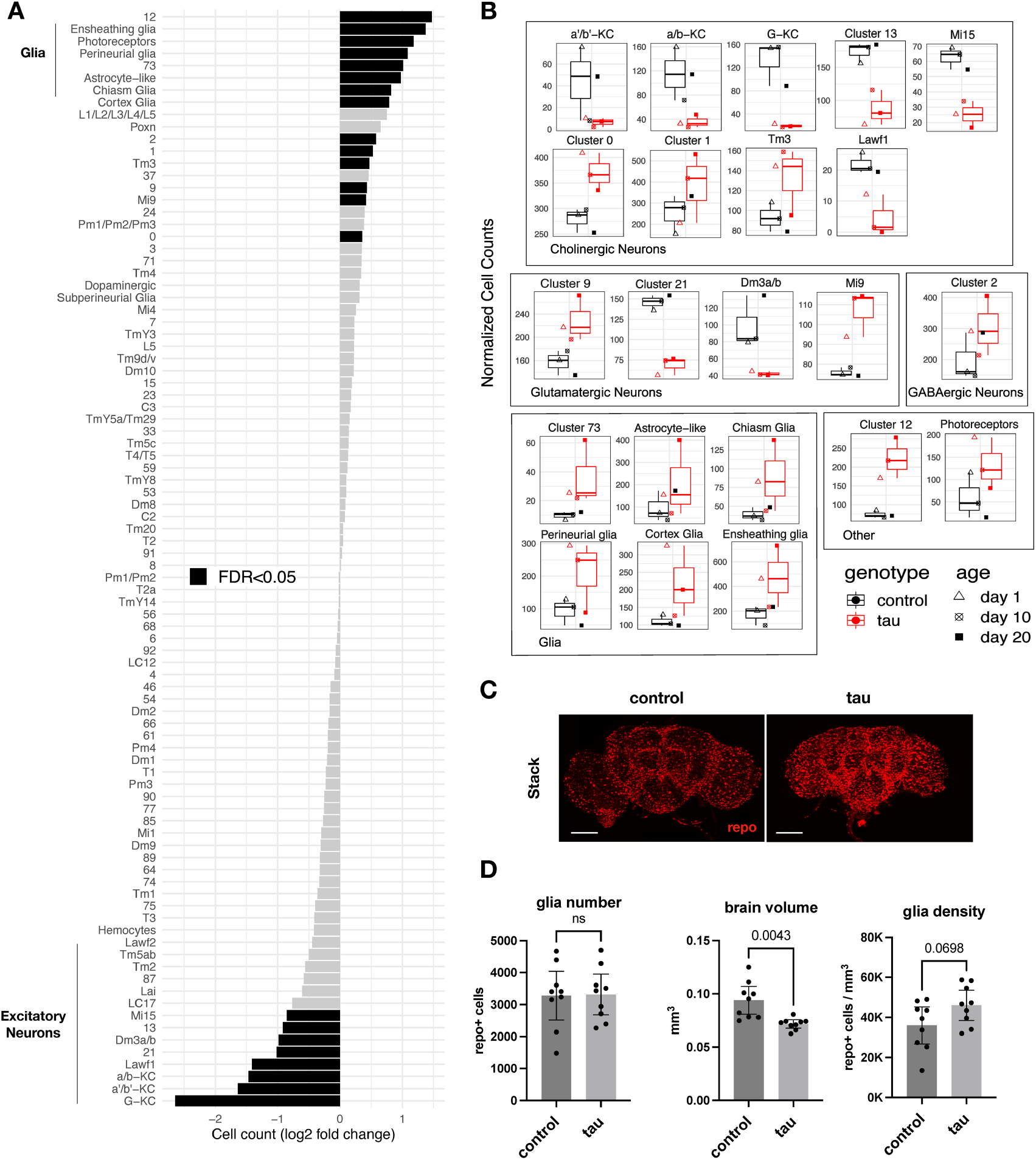
tau-triggered cell proportion changes in the adult brain. (A) Log2-fold change (log2FC) of normalized cell counts between *elav>tau^R406W^ (elav-GAL4 / +; UAS-tau^R406W^ / +*) and control (*elav-GAL4 / +*) animals. Cell clusters with statistically significant changes [False Discovery Rate (FDR) < 0.05] are highlighted in black. Many of these cell abundance changes were replicated in an independent dataset generated from 10-day-old animals (Figure S4). Since cell-type abundance estimates are relative between clusters, we also performed an adjusted analysis in which glia were assumed to be unchanged (Figure S6A). (B) Plots highlight cluster cell counts with significant differences between *elav>tau^R406W^* (red) and control (black) animals, including results for samples collected at 1- (triangle), 10- (cross-hatch square), or 20- (filled square) days. Box and whisker plots show minimum, median, and maximum values, along with lower and upper quartiles. See Figure S5 for complementary analysis based on deconvolution of bulk brain RNA-sequencing. (C) Whole-mount immunofluorescence of adult brains from 10-day-old flies. Glia are stained using the Anti-Repo antibody (red) in control (*elav-GAL4 / +*) and *elav>tau^R406W^* transgenic flies. Full Z-stack projection is shown. Scale bar = 100 microns. See also Figure S6B for additional immunostains for nuclei and actin. (D) Quantification of glia (Repo-positive puncta), brain volume, and glial density is shown. Error bars denote the 95% confidence interval. See also Figures S4-S6, Table S2.

Similar to our *Drosophila* tauopathy model, snRNAseq from postmortem human brain tissue has consistently suggested AD-associated increases in glial cell abundance, including astrocytes, oligodendrocytes, microglia, and endothelial cells (Lau et al., 2020; Zhou et al., 2020). However, one major limitation of both scRNAseq and snRNAseq analysis is that cell type abundance estimates are relative across the dataset. Therefore, a decline in neuronal subpopulations could lead to inflated abundance estimates of other, stable cell types. Indeed, whereas widespread neuronal loss is highly characteristic of AD (Davies and Maloney, 1976; Braak and Braak, 1991; Leng et al., 2021), systematic histopathologic studies in postmortem brain tissue do not support an absolute increase in microglia or astrocyte numbers, but rather a proportional increase in reactive glia in diseased tissues (Serrano-Pozo et al., 2013; Davies et al., 2017; Paasila et al., 2019). We therefore computed confidence intervals for cell abundance changes under an alternative model in which glia were assumed to be unchanging (Figure S6A). In this more conservative, adjusted analysis, only the neuroendocrine group (cluster 12) was increased and 15 excitatory neuronal subtypes were decreased.

In order to resolve the remaining ambiguity in potential glial cell changes, we performed immunofluorescence on whole-mount *Drosophila* brains (Figure 2C). Although the overall intensity of glial nuclear staining (anti-Repo) was increased in *elav>tau^R406W^* flies, quantification revealed no significant increase in absolute glial numbers. Instead, we found nominally increased glial density in tau animals after considering their reduced total brain volumes (Figure 2D). The increased intensity of antibody staining in tau brains may arise from enhanced antibody penetration, since similar changes are also seen for other markers (Figure S6B). Moreover, increased *repo* gene expression was not observed in either scRNAseq or in our previously published bulk-tissue RNAseq (Mangleburg et al., 2020). Overall, our results suggest that the apparent increase in glial cell abundance from scRNAseq is likely a consequence of proportional changes in single cell suspensions due to neuronal loss. While it is difficult to exclude more modest or selective regional changes, we conclude that similar to human postmortem tissue findings (Serrano-Pozo et al.; 2013), absolute glial numbers are largely stable following tau expression in the *Drosophila* brain.

### tau and aging exert cell-specific effects on brain gene expression

To our knowledge, the specific contributions of tau and aging on gene expression across heterogeneous cell types in the adult brain have not been systematically examined. In order to define the impact of aging on brain gene expression, we first quantified cell-specific transcriptional signatures in control flies (*elav-GAL4*) by performing differential expression analyses between the 3 timepoints from matched cell clusters (Figure 3A; Table S4). Overall, we define 5,998 unique, aging-induced differentially expressed genes. Based on gene ontology term enrichment, ribosome/protein translation and energy metabolism pathways were broadly dysregulated during aging, involving the majority of cell types (Table S5). We next used linear regression to examine tau-induced differential gene expression within each cell type, including adjustment for age as a covariate. Overall, a total of 5,280 unique genes were differentially expressed in at least one or more cell types (Figures 3B and S9A), and these results overlap significantly with our prior bulk RNA-seq in *elav>tau^R406W^* flies (Figure S7). Importantly, 93% of tau-induced differentially expressed genes were also triggered by aging in control flies. However, tau and aging appeared to have markedly distinct impacts when considering the distribution of gene perturbations across heterogeneous cell types (Figure 3C). Whereas aging broadly perturbed gene expression, tau-triggered changes were sharply polarized to excitatory neurons and glia. Moreover, tau-specific signatures predominated in selected cell types. For example, cholinergic Kenyon cells from the α’/β’ mushroom body lobes, which were among the most vulnerable cell types (Figure 2A), also had the greatest number of tau-induced gene perturbations (Figure 3B; Table S4). In fact, among 2,289 tau-induced differentially expressed genes within α’/β’ Kenyon cells, 2,139 (93%) were unique to tau and not similarly triggered in the corresponding cell type in aging control animals. We confirmed that the number of differentially expressed genes and affected cell types does not correspond to the spatial pattern of *MAPT* transgene panneuronal expression in the brain (Figure S8).

**Figure 3.**
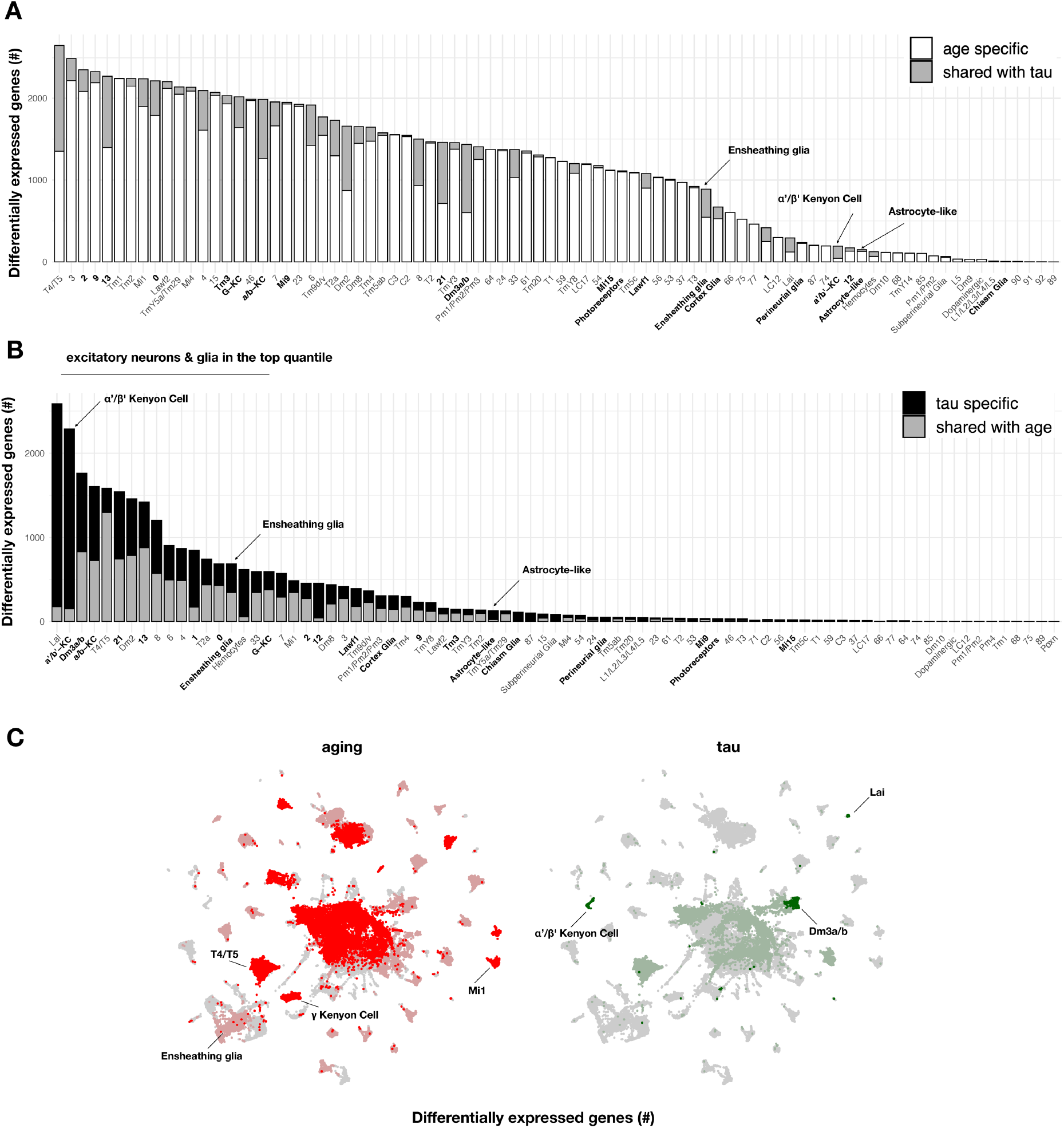
Aging-versus tau-triggered brain gene expression changes. (A) Aging has widespread transcriptional affects on most brain cell types. Number of aging-induced differentially-expressed genes (FDR<0.05) within each cell cluster is shown, based on comparisons of day 1 vs day 10 and day 10 vs day 20 in control animals only (*elav-GAL4 / +*). For each cell cluster, the number of gene expression changes unique to aging (white) or overlapping with tau-induced changes (gray) is highlighted. Labels for cell clusters with significant tau-induced cell abundance changes are shown in bold. (B) In contrast with aging, tau induces a more focal transcriptional response, with greater selectivity for excitatory neurons and glia. Number of tau-induced, differentially-expressed genes (FDR<0.05) within each cell cluster is shown, based on regression models including age as a covariate and considering both control and *elav>tau^R406W^* animals (*elav-GAL4 / +; UAS-tau^R406W^*/ +) at 1-, 10-, and 20-days. For each cell cluster, the number of gene expression changes unique to tau (black) or overlapping with aging-induced changes (gray) is highlighted. Labels for cell clusters with significant tau-induced cell abundance changes are shown in bold. Tau-induced gene expression changes from single-cell profiles significantly overlap with prior analyses conducted using bulk brain RNA-sequencing (Figure S7). (C) Uniform manifold approximation and projection (UMAP) plots show the number of aging- (red) versus tau- (green) triggered differentially expressed genes within each cell cluster. Color intensity represents the scaled number of differentially expressed genes. See also Figures S8-S9, Tables S4-S6.

Using functional enrichment analysis, we identify tau transcriptional signatures implicating altered inflammation, oxidative phosphorylation, and ribosomal gene expression (Figure S9B; Table S5). These pathways were prominently disrupted in excitatory neurons of the fly visual system, along with other central brain cholinergic and glutamatergic cell clusters. The pattern of transcriptional perturbation is also consistent with the established susceptibility of the mushroom body and optic lobes to tau-mediated neurodegeneration (Wittmann et al., 2001, Kosmidis et al., 2010). In other cases, we noted functional enrichments with greater specificity for selected cell clusters, such as altered signatures for mTOR signaling in glutamatergic cluster 21 and Foxo signaling in a subset of neuron types, including lamina intrinsic amacrine (Lai) cells and a cluster receptive to columnar motion (T4/T5). In addition, genes involved in mRNA splicing regulation were perturbed in another group of visual processing cells (T2a) as well as cholinergic cluster 7. Among non-neuronal cells, ensheathing glia, cortex glia, astrocyte-like glia, and hemocytes had the greatest number of tau-driven differential expression changes, highlighting signatures related to fatty acid metabolism and synaptic regulation. To further examine the robustness of our findings, we performed cross-sectional analyses, comparing our discovery dataset at day 10 with a larger replication dataset at the same age (26,000 cells vs 69,128 cells) (Table S6). Overall, there was a 90% overlap in tau-induced gene expression changes between the discovery and replication dataset, including significant overlaps in the most vulnerable excitatory neuron and glial cell clusters.

### tau triggers changes in neuronal innate immune signaling

Whereas most tau-induced genes strongly overlapped with aging, a minority overall were tau-specific (363 out of 5280 gene perturbations). Interestingly, this gene set was significantly enriched for mediators of the innate immune response, particularly NFκB signaling pathway components (Table S5). From *Drosophila* bulk brain RNA-seq data, we previously identified 7 gene coexpression modules perturbed by *tau^R406W^* expression using weighted correlation network analysis (Mangleburg et al., 2020). Among these, a 236-gene module was strongly enriched for innate immune response genes downstream of NFκB. In order to better understand the cell type-specific expression patterns, we next examined the innate immune coexpression module in our scRNAseq data. This immune signature was broadly detected in the adult fly brain, including both glia and many neuron types (Figures 4A and S10A). Moreover, expression of the immune module was strongly dysregulated by tau, with 50 out of 90 clusters showing significant changes (Figure 4B; Table S7). Tau activated the immune signature in the majority of affected cell types (86%, 43 out of 50 clusters). In particular, tau-triggered increases were noted in multiple excitatory neuron clusters (e.g., Dm3 glutamatergic cells in the visual system) as well as non-neuronal cells, including glia (e.g., ensheathing and cortex glia) and hemocytes. Conversely, in a selected subset of 7 clusters, tau attenuated expression of the innate immune module (Figure 4B), including excitatory neurons in the lamina and several Kenyon cell types that were among the most vulnerable to tau-triggered neuronal loss, based on cell abundance estimates (Figure 2A). Other tau-perturbed coexpression modules revealed distinct cell type-specific patterns (Figure S11). For example, a module enriched for synaptic regulators was markedly reduced in glia in response to tau, whereas expression was increased in multiple glutamatergic neuron subtypes.

**Figure 4.**
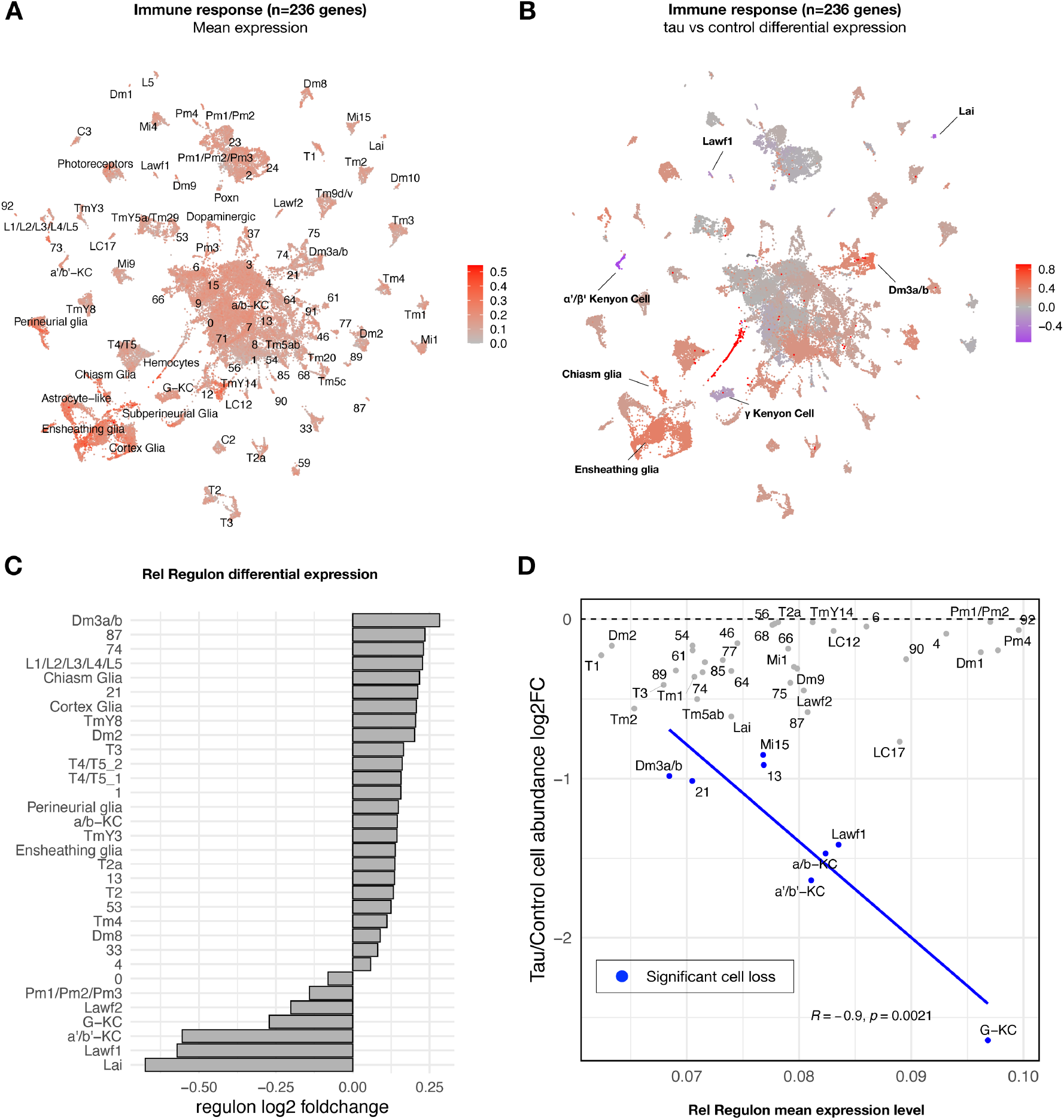
tau-induced changes in innate immune response genes and neuronal vulnerability. (A) Innate immune genes are expressed broadly in the adult fly brain, including both neurons and glia. Plot shows mean overall normalized expression by cell cluster among n=236 genes belonging to a tau-induced coexpression module that is significantly enriched for innate immune response pathways (Mangleburg et al., 2020). In this plot, gene expression was averaged across both *elav>tau^R406W^* and control cells; similar results are seen when stratifying by either age or genotype (Figure S10A). See also Figure S10D for experimental confirmation of NFκB/Rel protein expression in neurons and glia. (B) tau activates or suppresses innate immune response genes in a cell type-specific manner. Plot shows log2 fold-change mean expression per cell cluster for the same 236-gene immune response coexpression module, based on comparisons between *elav>tau^R406W^* (*elav-GAL4 / +; UAS-tau^R406W^* / +) and control (*elav-GAL4* / +) flies. See also Figure S10B and S10C for plots of curated NFκB signaling pathway genes and Figure S11 for similar analyses of other coexpression modules. (C) Log2 fold-change in Relish (Rel) regulon gene expression per cluster is shown, based on comparisons between *tau^R406W^* and control flies. All results were significant (FDR<0.05) based on regression models including age as a covariate. (D) Plot shows overall mean expression of the Rel-regulon (x-axis) versus tau-induced cell abundance change (y-axis). Among clusters with significant, tau-induced cell loss (denoted in blue, FDR<0.05; see also Figure 2A), cell abundance change was inversely correlated with Rel regulon expression (Pearson correlation: R = −0.9, p=0.0021). Many other cell types without significant cell abundance changes are also shown in gray. See also Figures S12, S13, and Tables S7-S10.

To confirm and extend our analysis of tau- and cell type-specific gene expression perturbations, we derived a complementary set of 183 transcription factor coexpression networks (regulons) based on our scRNAseq data. Specifically, regulons define coexpressed gene sets in which members are also predicted targets of a specific transcription factor (Van de Sande et al., 2020). Overall, clustering cells based on regulon enrichment recapitulates similar, expected relationships between annotated cell types (Figure S12; Table S8), and differential regulon analysis also revealed consistent tau-induced, cell type-specific transcriptional perturbations (Table S9). In particular, we examined the 442-gene regulon comprised of targets of the NFκB transcription factor ortholog in *Drosophila*, Relish (Rel), which is activated downstream of the *Drosophila* Imd (Immune deficiency) pathway, similar to the Tumor Necrosis Factor Receptor pathway in mammals (Myllymäki et al., 2014). The expression pattern of the Rel regulon and its differential expression in *tau* versus control flies were consistent with our findings for the immune coexpression module derived from bulk RNAseq, which includes both Imd, Rel, and multiple antimicrobial peptides that are activated by Rel (Figure 4C). We also obtained consistent results based on a manually-curated, 62-gene set including well established NFκB signaling pathway members (Figures S10B and S10C). We experimentally confirmed Rel expression in both neurons and glia in the adult fly brain using an available strain in which the endogenous protein harbors an amino-terminal GFP tag (Figure S10D).

### Expression signatures for neuronal vulnerability in tauopathy

In order to more directly model the relationship of transcriptional regulation and cellular vulnerability in tauopathy, we integrated regulon expression levels with cell abundance estimates from scRNAseq (Figure S13A). We hypothesized that innate immune signatures may be predictors of neuronal subtype vulnerability in tauopathy. We implemented regularized multiple regression in which cell-type specific regulon mean expression served as the predictor variable and tau-triggered cell abundance changes from scRNAseq provided the response variable. The analysis was restricted to cell clusters that show significant declines in *elav>tauR406W* flies. Out of 183 total regulons, Rel/NFκB activity was prioritized among 7 top predictors of vulnerability to tau-induced cell loss (Figures 4D and S13B). The Rel regulon remained a robust predictor in an expanded analysis including multiple technical variables as well as expression levels for an additional 2,793 curated functional pathways (Table S10). Importantly, for this analysis, regulon expression was averaged across both *elav>tau^R406W^* and control cells, rather than considering differential expression, and the vulnerable clusters include cell types in which Rel and its targets (Rel regulon) are either activated (e.g., Dm3) or suppressed (e.g, Gamma lobe of the Kenyon cells) in response to tau (Figure 4C). Interestingly, the inverse relationship with cell abundance is recapitulated when restricting consideration of Rel regulon activity in control animals, suggesting that basal NFκB signaling—in the absence of tau—may be a predictive marker for the severity of neurodegeneration (Figure S13C). In sum, our results identify NFκB targets and innate immune signaling as potential markers and/or mediators of vulnerability to tau-mediated neurodegeneration.

### Conservation of cell type-specific transcriptional signatures

To establish translational relevance, we next examined the conservation of cell type-specific transcriptional signatures between *Drosophila* and human brain. Using Pearson correlation and considering 5,630 conserved genes, we assessed correspondences between our *Drosophila* scRNAseq data and published snRNAseq from human dorsolateral prefrontal cortex (Mathys et al., 2019). Overall, inferred neuronal and glial cellular identities correlated well across species (Figures 5A and S14A). For example, a human microglial subcluster (Mic1) notable for association with high tau neuropathologic burden was correlated with the ensheathing glia cluster from *Drosophila*, indicating shared characteristic transcriptional signatures. Moreover, these 2 cell types showed significantly overlapping gene expression changes in response to AD or tau, respectively (hypergeometric test, p= 4.83×10^-5^) (Table S11). Crossspecies correlations in cell type-specific signatures were further replicated in an independent snRNAseq dataset from the human entorhinal cortex (Grubman et al., 2019) (Figure S14B).

**Figure 5.**
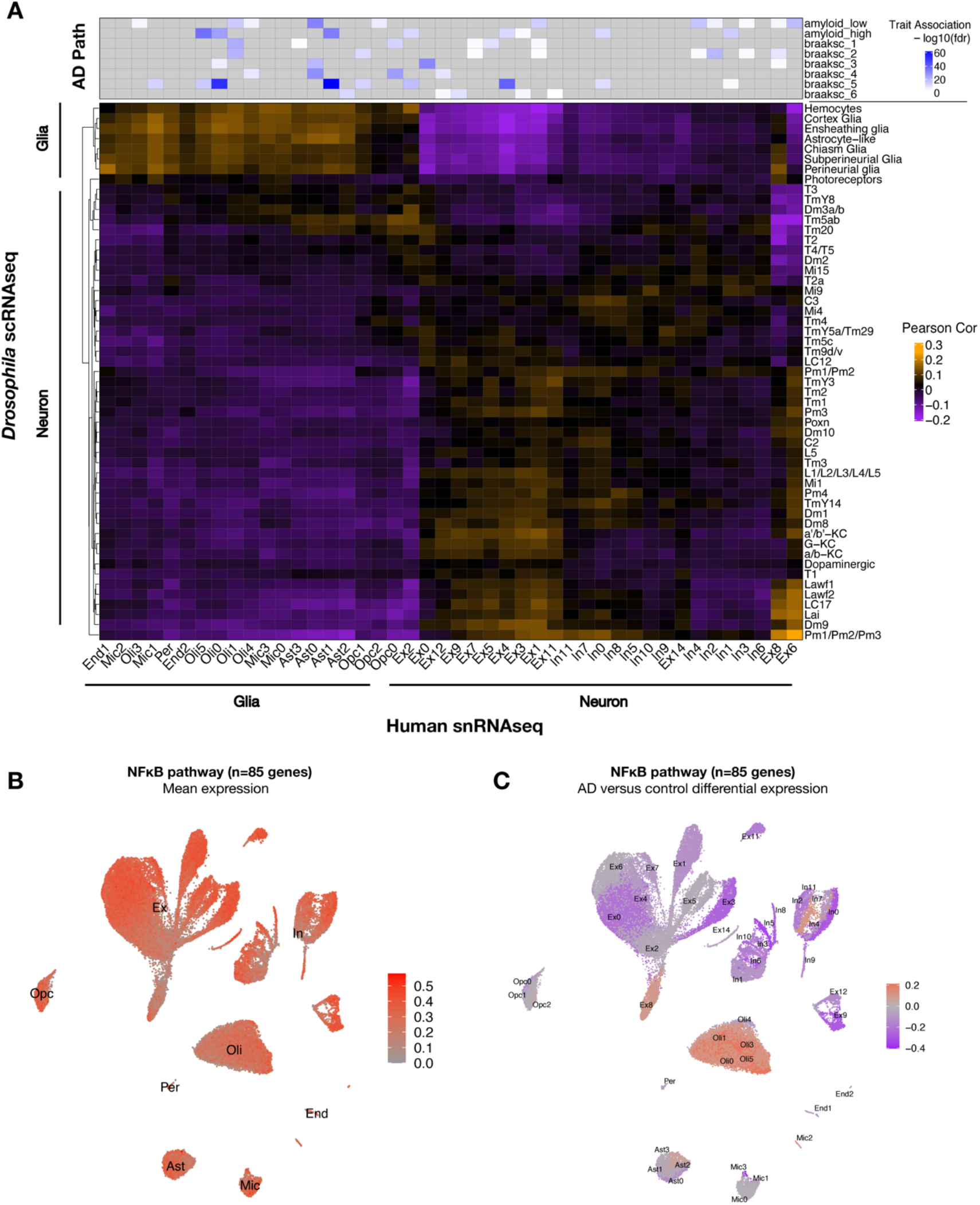
Conservation of cell type-specific gene expression signatures. (A) Heatmap shows Pearson correlation of gene expression (5,700 conserved, orthologous genes) between annotated cell clusters from *Drosophila* (rows) and human postmortem brain (column). Human brain single-nucleus RNA-sequencing (snRNAseq) was obtained from Mathys et al. (2019), including published cell-type associations with amyloid plaque burden and neurofibrillary tangle Braak staging (braaksc) (top). Annotated human cell types include endothelial cells (End), microglia (Mic), oligodendrocytes (Oli), pericytes (Per), astrocytes (Ast), oligodendrocyte precursor cells (Opc), excitatory neurons (Ex), and inhibitory neurons (In). (B) Innate immune mediators are expressed broadly in the human brain, including in neurons and glia. Plot shows mean expression by cell cluster for 85 human orthologs of NFκB signaling pathway members, based on reprocessing and analysis of the Mathys et al. snRNAseq data. (C) AD is associated with cell-type specific perturbation in NFκB signaling genes. Plot shows log2 fold-change mean expression per cell cluster for the same 85 NFκB signaling genes, based on comparisons of brains with Alzheimer’s disease (AD) pathology versus controls. See also Figure S14 and Table S11.

As introduced above, mediators of innate immunity are also highly conserved across species. Leveraging the published human snRNAseq data, we confirmed that NFκB signaling pathway genes are expressed across most cell types in human postmortem brain tissue, including both neurons and glia (Figure 5B). In the context of AD pathology, NFκB pathway gene expression appeared strongly down-regulated in most neurons from the dorsolateral prefrontal cortex, which are highly susceptible to degeneration, whereas expression was increased among oligodendrocytes, microglia, and astrocytes (Figure 5C). Interestingly, a subset of excitatory and inhibitory neuronal subclusters (Ex8 and In4, respectively) showed an AD-associated increase in expression. Thus, in both *Drosophila* and human brains AD pathology is associated with widespread changes in NFκB innate immune signaling, including either activation or attenuation in many distinct neuronal and non-neuronal subtypes.

## Discussion

Aging is the most important risk factor for AD, influencing both disease onset and progression. Based on longitudinal, single cell analysis in *Drosophila*, we discover that tau and aging activate strongly overlapping transcriptional responses: 93% of tau-induced differentially expressed genes are also perturbed by aging in control animals. Instead, tau and aging are distinguished by their spatial and cell type-specific impacts. Aging has a global influence on brain gene expression, affecting most brain cell types. By contrast, tau has a focal impact, polarizing the transcriptional response to a handful of cell types, including excitatory neurons and glia. The strong overlap between tau- and aging-induced gene expression signatures agrees with our prior analyses of bulk brain tissue (Mangleburg et al., 2020). We and others have also documented similar findings in AD mouse models, including both *MAPT* and *amyloid precursor protein* transgenics (Wan et al., 2020; Cummings et al., 2015; Gjoneska et al., 2015; Matarin et al., 2015; Hargis & Blalock, 2017). By contrast with animal models, cross-sectional studies of human postmortem tissue make it difficult to disambiguate the impact of aging from disease pathology on the brain transcriptome. However, our cross-species analyses highlight that most human brain cell types share transcriptional signatures with counterparts in the *Drosophila* brain. These correspondences comprise a cross-species atlas enabling studies of controlled experimental manipulations (*e.g.*, tau vs. aging) on homologous cell clusters between humans and flies.

Mechanistic dissection of cell type-specific vulnerability promises to reveal drivers for the earliest clinical manifestations of AD, such as the characteristic memory impairment accompanying the loss of excitatory neurons in hippocampus and associated limbic regions (Mrdjen et al., 2019; Fu et al., 2018). Given the transcriptional overlaps, one attractive model is that aging establishes a spatial pattern of vulnerable cell states that templates the subsequent tau-triggered neurodegeneration. However, as noted above, aging has wide-ranging impact across the brain and many cell types with robust aging-induced transcriptional responses in *Drosophila* are, in fact, resilient to tau-mediated neurodegeneration based on cell proportion changes (*e.g.*, clusters 2, 3 and T4/T5; Figures 2A and 3A). Moreover, the overlap between tau and aging does not reliably predict those cell types that are most vulnerable to neuronal loss (Figures 3A and 3B).

Differentially expressed genes triggered by tau and aging are nevertheless similarly enriched for many common biological pathways that may provide clues to cell type-specific mechanisms of vulnerability in neurodegeneration. Specifically, we document shared expression signatures for altered synaptic regulation, protein translation, lipid metabolism, and oxidative phosphorylation across heterogeneous cell populations, including excitatory neuron types that are particularly vulnerable to tau. Similar pathways have been implicated based on snRNAseq analyses from human postmortem brain (Mathys et al., 2019; Grubman et al., 2019; Lau et al. 2020) and several mouse AD models, including *MAPT* transgenics (Wang et al., 2022; Lee et al., 2021; Habib et al., 2020; Zhou et al., 2020).

Among the many dysregulated molecular processes, aging is characterized by a systemic pro-inflammatory state which has been called “immunosenescence” or “inflamma-aging” (Shaw et al., 2013; Hou et al., 2019). Genes encoding regulators of immunity, including *TREM2, CR1*, and many others, have been strongly implicated in AD susceptibility by human genetics (Bellenguez et al., 2022), and abundant evidence now supports a key role for many such genes among glial cells (Wang et al., 2015; Zhou et al., 2020; Keren-Shaul et al., 2017). We previously identified an age-associated *Drosophila* innate immune response signature that is amplified by tau (Mangleburg et al., 2020). Here, we significantly extend these observations, leveraging the cellular resolution afforded by single cell profiles. First, we discover that this immune coexpression module, including many NFκB / Rel signaling factors and targets, is broadly expressed in the adult fly brain, including both neurons and glia, and we confirm similar findings in snRNAseq data from human postmortem brain. Second, we show that tau can either activate or attenuate NFκB immune pathways in a cell type-specific manner, with tau-triggered decreases in expression apparent in neurons with the greatest proportional cell loss. Lastly, models integrating cell type-specific gene transcriptional expression and cell abundance changes suggest that basal Imd signaling strength (i.e., Rel regulon activity) predicts the severity of tau-triggered neuronal decline. Overall, our results suggest that besides the well-established requirements in glia (see below), innate immune response pathways may also have important, cell autonomous roles in modulating neuronal vulnerability to tau pathology in AD. Indeed, both insect and mammalian neurons express evolutionary-conserved innate immune signaling pathways, including Toll-like receptors and NFκB signal transduction components, and these pathways can be triggered by infection or other cellular insults (Lehnardt et al., 2003; Tang et al., 2007; Cao et al., 2013; Cho et al., 2013; Welch et al., 2022). In addition, NFκB immune signaling pathways have been coopted for diverse, non-canonical functions, such as in neurodevelopment and synaptic plasticity (Okun et al., 2011; Gutierrez and Davies, 2011; Nguyen et al., 2020). Knockdown of *Rel* in *Drosophila* neurons promotes survival (Kounatidis et al., 2017), whereas activation of the Rel signaling pathway leads to neurodegeneration (Cao et al., 2013). Consistent with our results, a recent reanalysis of snRNAseq data from Mathys et al. (2019) revealed AD-associated perturbation of NFκB immune pathways in excitatory neurons, possibly triggered by DNA double strand breaks (Welch et al., 2022).

Our scRNAseq analyses also highlight a robust, tau-induced transcriptional response among *Drosophila* glia. This result is consistent with several brain gene expression studies from both humans and mouse models that strongly implicate altered transcriptional states and/or increased numbers of AD-associated glial subtypes, including oligodendrocytes, astrocytes, and microglia (Mathys et al., 2019; Grubman et al., 2019; Lau et al., 2020; Zhou et al., 2020; Habib et al., 2020). Although our analyses initially suggested a possible tau-triggered increase in glial abundance in the brain, on direct examination, we documented stable absolute numbers but increased density of glia due to brain atrophy. Systematic histopathologic studies in human brain tissue have similarly revealed predominantly reactive changes with overall stable numbers of both astrocytes and microglia (Serrano-Pozo et al., 2013). We conclude that potential increases in disease-associated glia inferred exclusively from single cell profiles should be interpreted cautiously, and additional experimental investigations may ultimately be required to resolve whether they result from (i) absolute changes in cell number, (ii) activation and/or transformation of cell states, (iii) or proportional changes due to primary perturbations in other brain cell types. Nevertheless, glial-specific experimental manipulations of immune regulators in many experimental models, including NFκB and the AD susceptibility gene *TREM2*, can potently modify neurodegeneration, consistent with cell non-autonomous requirements (Walter, 2016; Kounatidis et al., 2017; Petersen et al., 2012; Hakim-Mishnaevski et al., 2019; Fuhrmann et al., 2010; Town et al., 2008; Leyns and Holtzman, 2017; Wang et al., 2022). By contrast with mammals, glia represent only 5-10% of all cells in the *Drosophila* brain (Ito et al., 1995; Schmidt et al., 1997; Awasaki et al., 2008). Nevertheless, *Drosophila* glial subtypes recapitulate the diversity of functions and morphologies of mammalian glia (Doherty et al., 2009; Freeman, 2015; Kremer et al., 2017; Stork et al., 2012). Although the myeloid hematopoetic lineage is not present in flies, which therefore lack microglia, ensheathing glia can similarly respond to cellular injury and scavenge debris (Doherty et al., 2009). Indeed, our cross-species analysis demonstrates shared transcriptional signatures between corresponding glial subtypes, consistent with our findings of conserved responses to tau-mediated neuronal injury.

## Acknowledgements

We are grateful to Pinghan Zhao, Tom Lee, Alma Perez, Katy Zhu, Akash Tarkunde and Bismark Amoh for assistance with *Drosophila* brain dissections. We also thank the Bloomington Drosophila Stock Center, the Vienna Drosophila RNAi Center, FlyBase (Gramates et al., 2017), the Developmental Studies Hybridoma Bank, and Biorender. This study was supported by grants from the National Institutes of Health (NIH) (R01AG057339, R01AG053960, U01AG061357, and U01AG046161). In addition, we utilized BCM core resources, including the Intellectual and Developmental Disabilities Research Center (P50HD103555), Genomic and RNA Profiling (S10OD023469), and Single Cell Genomics (S10OD025240 and Cancer Prevention Research Institute of Texas grant RP200504). H.Y. is additionally supported by the Parkinson’s Foundation (PF-PRF-830012) and the Alzheimer’s Association (AARF-21-848017). J.M.S. was additionally supported by the Huffington Foundation, McGee Family Foundation, the Jan and Dan Duncan Neurological Research Institute at Texas Children’s Hospital, and The Effie Marie Caine Endowed Chair for Alzheimer’s Research.

## Author contributions

Conceptualization, TW, ZL, IA-L, JB, JMS; Investigation, TW, JMD, HY, CG, JD, BTP, RA-O; Formal Analysis, TW, JMD, BTP, RA-O; Writing – Original Draft, TW, JMS; Writing – Review and Editing, TW, JMD, HY, CG, JD, BTP, RA-O, ZL, IA-R, JB, JMS; Funding Acquisition, JB, ZL, JMS; Supervision, JB, ZL, IA-R, JMS.

## STAR METHODS

### RESOURCE AVAILABILITY

#### Lead contact

Further information and requests for resources should be directed to and will be fulfilled by lead contact, Joshua M. Shulman.

#### Materials availability

This study did not generate any new unique reagents.

#### Data and code availability

- All original single cell sequencing data have been uploaded to the Accelerating Medicines Parternship (AMP)-AD Knowledge Portal on Synapse and can be accessed through the DOI: https://doi.org/10.7303/syn35798807.1. All raw data required for analytic replication have been uploaded. Full resolution immunofluorescence images will be shared by lead contact upon request.
- This paper does not report original code.
- Any additional information required to reanalyze the data reported in this paper is available from the lead contact upon request.

### EXPERIMENTAL MODEL AND SUBJECT DETAILS

#### Human subjects

No new data from human subjects were generated for this study. Previously published, available snRNAseq data from human postmortem brain were obtained from Mathys et al. (2019) and Grubman et al. (2019) in order to evaluate cross-species correspondences in cell type-specific expression signatures. The Mathys data is comprised of snRNAseq from the dorsolateral prefrontal cortex (DLPFC) from 48 brain autopsies with varying AD neuropathology (amyloid plaque and tau neurofibrillary tangle burden), including 24 with no significant pathology (controls) and 24 cases with mild to severe AD pathology. Subjects were balanced for sex (12 male and 12 female), and age (median age at death = 87 for both groups). The Grubman data is comprised of snRNAseq from the entorhinal cortex of 12 brain autopsies, including 6 AD pathologic cases and 6 controls without significant AD pathology. Subjects in the Grubman data were also age-matched, with a median age of 83 and 80 for the AD case and control groups, respectively.

#### *Drosophila* stocks

For scRNAseq libraries generated in this study, *w^1118^; UAS-tau^R406W^* flies (0N4R isoform, 383 amino acids), described in (Wittmann et al., 2001; Mangleburg et al., 2020), were crossed with the pan-neuronal driver *elav^C155^-Gal4*, producing the experimental genotypes: *elav-Gal4/ +;UAS-tau^R406W^/ +* or *elav-Gal4 / Y; UAS-tau^R406W^ / +*. Controls were generated by outcrossing *elav-Gal4* with *w^1118^* animals, producing *elav-Gal4 / +* or *elav-Gal4 / Y*. Adult progeny from experimental crosses were subsequently aged to 1-, 10-, or 20-days for dissection and library generation. Flies were raised on standard molasses-based media at 25°C in ambient lighting. We also utilized a *Rel-GFP* strain (*y, w; PBac{GFP.FPTB-Rel}VK00037*), which is an endogenous protein trap allele, encoding a fusion protein with GFP at the Rel amino-terminus.

### METHOD DETAILS

#### *Drosophila* brain dissociation

For scRNAseq profiling of *elav>tau^R406W^* and control flies, 16-18 dissected and intact *Drosophila* brains were combined and dissociated for each experimental condition (6 total samples: 2 genotypes x 3 timepoints). An equal number of male and female animals were combined for each condition. For the replication dataset, triplicate samples for the identical *elav>tau^R406W^* and control genotypes were prepared at day 10 (6 total samples). Adult fly brains were dissected out of the cuticle using sharp forceps in 1X PBS and dissociated following published protocols (Davie et al., 2018). Dissected brains in solution were first centrifuged at 800g for 3 min, resuspended, and dissociated by incubating with 50uL of dispase (3mg/mL, Sigma) and 75uL of collagenase I (100mg/mL, Invitrogen) for 2 hours at 25°C while shaking at 500 RPM. Cell suspensions were mixed by gentle pipetting 3-4 times every 5 minutes in the first hour, and every 10 minutes in the second hour. Resulting cell suspensions were pelleted by centrifugation at 400g for 5 min at 4°C, washed in 1000uL ice cold PBS, pelleted, and resuspended in 400uL ice cold PBS with 0.04% bovine serum albumin. Cell suspensions were passed through a 10um pluriStrainer cell strainer (pluriSelect) to ensure that undissociated tissue were removed and a single cell suspension was obtained. Cell concentration and viability were assessed using a hemocytometer under a fluorescent microscope after staining with NucBlue and Propidium iodide (Invitrogen). Fresh, intact single cell suspensions were immediately used for single cell library preparation.

#### Single cell library preparation and sequencing

Single cell libraries were prepared per manufacturer’s protocol for the Chromium Single Cell Gene Expression 3’ v3.1 kit (10x Genomics) by the BCM Single Cell Genomics Core. 16,000 cells were added to each channel with a target recovery rate of 10,000 cells per library. Cells, reverse transcription (RT) reagents, gel beads containing barcoded oligonucleotides, and oil were loaded on a Chromium controller (10x Genomics) to generate single cell GEMs (Gel Bead-In-Emulsions) where full length cDNA was synthesized and barcoded for each individual cell. GEMs were subsequently broken and cDNAs from each single cell were pooled. Following clean up using Dynabeads MyOne Silane Beads (Invitrogen), cDNA was amplified by PCR. The amplified product was fragmented to optimal size before end-repair, A-tailing, and adaptor ligation. Final library was generated by amplification. Completed libraries were sequenced using the Baylor Genomic and RNA Profiling Core on the Illumina NovaSeq 6000 platform with a minimum depth of 300,000,000 reads per sample (on average 463M reads per sample). A total of 12 high-quality libraries were generated (6 libraries for the discovery and replication datasets, respectively). Illumina BCL files were demultiplexed into FASTQ files by calling the Cell Ranger 4.0.0 *mkfastq* function. FASTQ files were aligned to the *Drosophila* reference genome (BDGP6.22.98) and quantified using the Cell Ranger 4.0.0 *count* pipeline. The human microtubule associated protein tau (MAPT) mRNA sequence (isoform 3, NCBI Reference Sequence NM_016834.5:151-1302) was appended to the *Drosophila* reference genome for assessing MAPT transgene expression levels. Given the 10x recovery rate estimations, the cell calling algorithm in Cell Ranger was applied by setting the -- expect-cells parameter in *count* to 10,000 for each library, thus filtering out partitions that likely did not contain single cells. Filtered count matrices were loaded into Seurat v3 in R for additional quality control and downstream analyses. Cells were removed from the data object if the number of unique genes per cell were less than 200 or greater than 3,000, or if the proportion of mitochondrial reads per cell was greater than 20%. Filtered count matrices from Cell Ranger are available to download with the *Drosophila* scRNAseq data on the Synapse AMP-AD Knowledge Portal.

#### Normalization, integration, and clustering

Gene expression was first normalized independently per library using a regularized negative binomial regression approach as implemented by SCTransform (Hafemeister & Satija, 2019). 5,000 highly variable features (HVG) were used for normalization while accounting for percent mitochondrial reads. Variable features were defined and ranked by computing the variance of standardized gene counts after loess-based adjustment of mean-variance relationships (Stuart et al., 2019). Residuals of the fitted regression models were used as normalized gene expression values for HVGs. All libraries normalized via SCTransform were integrated using the canonical correlation analysis (CCA) pipeline in Seruat v3 to correct for batch effects and facilitate identification of similar cell identities across conditions. Highly ranked HVGs shared across all libraries were used as integration features. Integration anchors across libraries (correspondences of the selected features between cells) were computed over the first 30 CCA dimensions in the combined dataset and then used to inform the subsequent integration and grouping of cells. After integration, Seurat v3 was used for principal component analysis (PCA) and cell clustering. 100 principal components of the integrated dataset were used for graph-based clustering and Louvain algorithm optimization as implemented in FindNeighbors and FindClusters. The final resolution in FindClusters was set to resolution = 2, yielding 96 cell clusters in our dataset. We selected this resolution to replicate the clustering pattern of a similarly processed *Drosophila* whole brain scRNA-seq dataset (Davie et al., 2018). 100 PCs were used to embed cells in 2D space via uniform manifold approximation and projection (UMAP).

Normalization of gene counts used in differential expression analysis, cell cluster marker gene computation, cell identity annotation, and other applications directly comparing gene expression levels between cell clusters were computed separately on the non-integrated gene expression data using the NormalizeData function in Seurat v3. In brief, for each gene in each cell, unique molecular identifiers (UMI) were divided by the sum UMIs in that cell, multiplied by a scalar (10,000), and log transformed. However, cell cluster membership (clusters 0-95) was defined using the integrated dataset as described above. The 6 additional libraries that comprise the day 10 replication dataset were clustered, integrated, and analyzed separately using the identical pipeline.

#### Doublet detection

DoubletFinder was applied per library to predict and remove heterotypic doublets, leaving a total of 48,111 high quality single cells in the discovery dataset. For each library, artificial doublets were generated from the existing data. Principal component analysis (PCA) was performed after merging the real and artificial data and a distance matrix was generated with the first 40 PCs to compute the proportion of artificial K-nearest-neighbors (pANN) for each cell. PC neighborhood size (pK) for computing pANN was estimated for each library as previously described (McGinnis et al., 2019). The number of suspected doublets per library was estimated and cells were ranked by pANN for removal. Total doublet proportion *Y* for each library was computed based on a custom linear equation of the input-to-multiplet estimation provided by the 10x Chromium documentation: *Y* = 5.272 * 10^-4^ + 7.589 * 10^−6^ x, *x* being the number of recovered intact cells after the initial filtering criteria described above. The linear equation was generated based on recovery estimations in the manufacturer’s protocol. Adjustment of the estimated doublet proportion for undetectable homotypic doublets was applied in DoubletFinder by using the Seurat clustering classifications at resolution = 2 as described above.

#### SCENIC regulons

Gene regulatory networks (regulons) were computed using the python implementation of SCENIC (pySCENIC). Raw gene abundances (UMIs) for 48,111 high quality cells were exported as a loom object via loompy, and pySCENIC was implemented as described in Van de Sande et al. (2020). Putative gene targets for the published list of 815 *Drosophila* transcription factors (TFs) (see Key Resources Table) were inferred by tree-based regression (GRNBoost2) where expression of each gene was regressed on TFs, producing a list of adjacencies connecting TFs to their target genes (non-mutually exclusive). In the cisTarget step, modules were retained for further analysis if the regulatory motif of its parent TF was enriched among most gene members. Within retained modules, genes lacking enrichment of the appropriate motif were pruned. TF-motif annotations and pre-computed motif-gene rankings were obtained from https://resources.aertslab.org/cistarget/, *Drosophila* v8; motif search space encompassed up to 5Kb upstream of transcription start sites and intronic regions. This pipeline identified 183 regulons, encompassing 7,134 out of 14,907 genes in the transcriptome dataset (Table S8), and cell-level activity for each regulon was computed by a ranking and recovery approach using pySCENIC AUCell. Within each cell, genes were ranked by expression level in a descending order, then the cumulative number of genes recovered belonging to a regulon at each rank was recorded. An area under the curve (AUC) was calculated after applying a default cutoff at the 95^th^ percentile of gene ranks, and is used to infer regulon activity. High AUC scores indicate greater representation of a given regulon among the top 5% of highly expressed genes in a cell. AUC scores for the 183 regulons across 48,111 cells were used for unsupervised clustering by UMAP for visualization of cell relationships based on gene regulatory networks (Figure S12A).

#### Cell identity annotation

We searched for cell identities of the 96 defined clusters by consolidating a series of 4 analytic approaches (Figure S2A). Two published datasets were used as references for our annotation procedure, including 56,902 cells from adult wildtype *Drosophila (w^1118^* and *DGRP-551*) brains profiled at days 0, 1, 3, 6, 9, 15, 30, and 50 (Davie et al., 2018), as well as 109,743 cells from adult Canton-S *Drosophila* optic lobes at day 3 (Özel et al., 2020). Cell clusters in these references were previously annotated using available literature-based cell markers or statistical inference with published bulk RNA-sequencing of reporter-targeted cell types. The Davie et al. dataset contained 87 cell clusters (Seurat FindClusters res=2.0) with 41 assigned cell identities. The Özel et al. (2020) dataset was clustered at a higher resolution (Seurat FindClusters res=10), containing 200 cell clusters and 87 assigned cell identities. First, Scmap-cluster was used to compute gene expression correlation between each cell in our dataset to all defined clusters in the Davie and Özel datasets. 500 genes with higher-than-expected dropouts were selected as correlation features as described in (Kiselev et al., 2018). The cosine similarity, Spearman and Pearson correlations of these features were subsequently computed between each cell in our dataset and all reference cluster centroids. For a cell to be mapped to a reference cluster, 2 out of 3 similarity scores must be concordant, and at least 1 must be greater than 0.7. Second, we intersected the top 20 cluster markers for each cell cluster (ranked by log2 fold change) in our dataset with the top 20 markers in each reference cluster. Cluster markers (cluster-enriched genes) for our 96 cell clusters were computed by differential expression analysis of the non-integrated, normalized gene abundances, comparing each cell cluster against all remaining cells. Markers were defined as positively differentially expressed genes (log2 fold-change greater than 0.1, BH-corrected p-value < 0.05) when comparing cells in given cluster versus all remaining cells in the dataset. Cell clusters were “mapped” to a reference cluster in the Davie dataset (whole brain reference) if at least 13/20 top markers were shared. Likewise, a cluster was mapped in the Özel dataset (optic lobe reference) if at least 7/20 top markers were shared. These cutoffs were empirically determined by maximizing the number of best matches. Cell cluster markers for our dataset are listed in Table S3. Third, a trained neural network classifier for adult neurons as described in Özel et al. (2020) was implemented in python to label optic lobe neurons in our dataset. Log-normalized expression of 533 genes (out of the 587 genes in the Özel adult training set) across all cells were used as the input for the classifier. Finally, we checked for positive expression of well-established cell markers (Table S12) in each cluster as shown in Figure S2B and S3. Most cell cluster annotations were evaluated and consolidated based on best agreement across two or more approaches within or across the Davie and Özel references. Less certain annotations were visually inspected in UMAP space to check for proximity with adjacent clusters and manually evaluated for cell marker expression. Cell-level confidence for scmap assignment (similarity score) or neural network classifications (confidence score) were also manually evaluated. Results from the Özel reference was prioritized for optic lobe neurons, especially for cell clusters that may be heterogenous in the Davie reference (Dm8/Tm5c, TmY14, Tm9, Tm5ab, Mt1). Pm neurons, chiasm glia, and subperineurial glia did not reach consensus across 2 or more approaches and were thus deemed less confident annotations. Several other optic lobe cell types were well mapped in a single approach to the Özel et al. (2020) dataset (TmY8, TmY3, Tm5c, Tm5ab, Tm20, Dm2, Dm8, Mi9, LC12, and LC17), where robust metrics were observed from the optic lobe neural network predictor or with scmap. Confirming our cell identity correspondences with the published scRNAseq datasets, we found high correlation among normalized gene expression when comparing individual cells in our dataset with the cluster-level means of the transcriptome in reference clusters as computed by cosine similarity (Figures S2C and S2D). The cosine similarity score between each annotated cell in our dataset and its cluster-level counterpart in the Davie or Özel references were computed based on shared non-dropout (count > 0) genes, i.e., the transcriptome of each cell in our data was correlated to the cluster-level mean of corresponding genes in a reference cluster. Lower similarity scores may reflect gene expression changes induced by tau pathology, less confident annotation (in this study or in the references used), clustering resolution differences, or high variance in the reference cluster.

To annotate the replication scRNAseq data (69,128 cells), labels from the completed dataset above (48,111 cells) were transferred using the Seurat v3 FindTransferAnchors and TransferData functions. In brief, pairs of similar cells between the reference and query dataset were identified using a mutual nearest neighbor approach after projecting the replication dataset onto the reference dataset in PCA-reduced space. The 5,000 most variable genes in the new dataset were used for the dimensional reduction. Each cell was assigned a score and a predicted label from the reference dataset. Cell level metadata for both the discovery and replication datasets are uploaded to the Synapse AMP-AD Knowledge Portal as noted in the key resource table.

#### Cell abundance quantification

After annotation, cell counts for each assigned cluster (90 clusters) were first quantified per library (6 libraries, ages: day 1, 10, 20; genotypes: control, tau), and treated as count data. To adjust for extreme proportional differences in cell composition across libraries and differences in the total number of cells captured per library, cell counts were normalized using negative binomial generalized linear models (NB-GLM) as implemented in DESeq2 (Love et al., 2014). In brief, raw cell counts were modeled using NB-GLM with a fitted mean and a cluster-specific dispersion factor. Dispersion factors were computed based on mean count values using an empirical Bayes approach as described in Love et al., (2014). The fitted mean is composed of a library-specific size factor and a parameter proportional to the true counts in each cluster per library. To compute size factors per library, raw counts were organized in a matrix such that rows represent clusters and columns represent samples (libraries). Raw counts were first divided by the row-wise geometric means and then divided by the per-column median of resulting quotients (size factor) to obtain normalized cell count values per cluster. These normalized cell counts were used to generate the plots in Figure 2B. The 3 age groups for each genotype (day 1, 10, 20) were combined to produce an n=3 comparison of cell counts between tau and control animals.

#### Deconvolution of fly RNA-sequencing data

Deconvolving bulk-tissue RNA-sequencing data into estimated proportions of cell populations was performed by implementing Multi-subject Single-cell Deconvolution (Wang et al., 2019) using default parameters. MuSiC leverages cell-specific expression data from annotated scRNA-seq datasets and weighted non-negative least squares regression to characterize cell compositions of bulk tissue gene expression data. This approach accounted for gene expression variability across samples and cells, thus upweighting the most consistently expressed genes across samples or cells for deconvolution. Whole-head RNA-sequencing counts of experimental conditions identical to those in this study were taken from Mangleburg et al. (2020) and used as input for deconvolution. Specifically, cell counts for n=2 control and n=3 tau^R406W^ samples at days 1, 10, and 20 were deconvolved (15 samples total, each sample is a homogenate of 100 heads). 56,902 cells from a published *Drosophila* whole brain scRNAseq dataset was used as an orthogonal reference for deconvolution, providing cell specific transcriptional profiles from wildtype control animals (*w^1118^ & DGRP-551*) (Davie et al., 2018). Individual scRNA libraries were treated as subjects in the MuSiC pipeline for evaluating gene expression variability in marker gene weighting. Both annotated and unannotated cell clusters in the reference scRNAseq dataset were included in the deconvolution pipeline. Select cell clusters with non-zero estimated proportions across 2 or more timepoints were plotted in Figure S5.

#### Immunofluorescence and confocal microscopy

10-day-old female controls (*elav-GAL4* / +) or *elav>tau^R406W^ (elav-GAL4 / +; UAS-tau^R406W^*/ +) were used for glial quantification immunofluorescence experiments. The animals were anesthetized with CO2 and brains were dissected with forceps and fixed in 4% paraformaldehyde (PFA) overnight at 4°C. After fixation, PFA was aspirated and replaced with PBS with 2% Triton-X (PBST) and incubated at 4°C overnight for tissue penetration. Residual air trapped in brain tissues were removed by placing samples under a vacuum for 1 hour at room temperature. The brains were then incubated in blocking solution (5% normal goat serum in PBST) at room temperature, rocking for 1 hour. Primary antibodies were diluted in 0.3% PBST and samples were incubated in primary at 4°C, rocking for at least 24 hours. The primary antibody solution was aspirated, and the samples were washed with PBST (two quick washes followed by three 15-minute washes). Samples were incubated in secondary antibodies at room temperature, rocking for 2 hours. The secondary antibody solution was then aspirated and the samples were washed with PBST (two quick washes followed by three 15-minute washes). DAPI stain, when applicable, was added in the secondary antibody step. Whole brains were then mounted in Vectashield® antifade mounting medium (Vector Laboratories, H-1000-10) and stored in the dark at 4°C until imaged. Samples were imaged on a Leica Microsystems SP8X confocal microscope. Z-stacks covered the entirety of whole mount brains. We used the following antibodies and dilutions: mouse anti-Repo (8D12, 1:500 for glial quantification experiment, 1:50 for Rel experiment, DSHB), rat anti-Elav (7E8A10, 1:100, DSHB); rabbit anti-GFP (1:500; GeneTex), conjugated A488-Phalloidin antibody (1:500; ThermoFisher), Cy^TM^3 AffiniPure Goat Anti-Mouse (H+L) (1:500; Jackson ImmunoResearch Laboratories), Alexa 647-conjugated goat anti-Rabbit IgG (1:500; Jackson ImmunoResearch), Alexa Fluor 488 Donkey anti-Mouse IgG (H+L) (1:500; Jackson ImmunoResearch), Cy^TM^3 AffiniPure Goat Anti-Rat IgG (H+L) (1:500; Jackson ImmunoResearch).

#### Bulk-tissue RNA-sequencing data

Bulk-RNA sequencing data and weighted correlation network analysis (WGCNA) co-expression modules of the experimental conditions described in this study were obtained from Mangleburg et al. (2020). WGCNA module expression activity in scRNAseq was computed by taking the mean of module member genes within each cell. Cluster-level expression activity of WGCNA modules was then estimated by averaging the cell-level activity across all cells in a given cluster, and was subsequently used to compute the log2 fold change of tau versus control expression activity.

#### *Drosophila* NFκB signaling mediators

A list of *Drosophila* NFκB signaling pathway members was generated based on manual curation from published studies (Valanne et al., 2011, Myllymäki et al., 2014, Kounatidis et al., 2017, and Li et al., 2020), and included the following genes: *PGRP-LE, imd, Tak1, key, Rel, eff, PGRP-LC, bsk, akirin, Jra, sick, Tab2, IKKbeta, Uev1A, ben, Dredd, Fadd, PGRP-LA, Diap2, Diap1, egr, Traf6, trbd, pirk, casp, PGRP-LB, PGRP-LF, dnr1, scny, RYBP, PGRP-SC1a, PGRP-SC1b, PGRP-SC2, CYLD, POSH, spirit, spheroide, spz, PGRP-SA, PGRP-SD, pll, Myd88, dl, Gprk2, Deaf1, Tl, psh, grass, modSP, Dif, mop, tub, cact, nec, Pli, 18w, Toll-4, Tehao, Toll-6, Toll-7, Tollo*, and *Toll-9.*

### QUANTIFICATION AND STATISTICAL ANALYSIS

#### Statistical testing of normalized cell abundance

Statistical testing of the log2 fold change (log2FC) of tau versus control normalized cell abundance was performed using negative binomial-generalized linear models (NB-GLM) as implemented in DESeq2. Age was treated as a covariate, and a Wald test was performed on the coefficient of the genotype variable using the following model: log2(cell count) ~ age + genotype. Using DESeq2, log2FC was computed for each cluster (*elav>tau^R406W^* vs control) based on maximum-likelihood estimation after fitting the GLM. Raw log2FC values were transformed using an adaptive shrinkage estimator from the “ashr” R package as implemented in DESeq2 to account for clusters with high dispersion or low counts. These transformed log2FC values were then used for cell abundance analysis and interpretation. A Benjamini-Hochberg (BH)-adjusted p-value < 0.05 was used to establish significance of Wald test statistic. In order to generate the plots for Figure 2B, normalized cell counts were obtained using the *counts* function in DESeq2. Results were visualized using box and whisker plots, including the following values: median, minimum / maximum, and lower / upper quartiles.

In order to better understand how relative changes might influence cell abundance estimates, we inferred confidence intervals for cell cluster log2FC values (Figure S6A). Based on experimental ground truth (Serrano-Pozo et al., 2013), the log2FC value for 7 cell clusters (Ensheathing glia, Perineurial glia, Astrocyte-like glia, Cortex glia, Chiasm glia, Subperineurial glia, and Hemocytes) was centered to zero. Specifically, the value of each glial cluster was iteratively subtracted from the log2FC values for all other clusters, establishing a minimum and maximum log2FC value for all cell clusters. We predict that the true cell abundance falls within this computed range, after accounting for potential proportional influences. A range that includes zero thus suggests there may be no true change between tau and control.

#### Repo quantification

Quantification of glia from confocal immunofluorescence digital microscopy was performed using Imaris (v9.9.1) imaging software. We counted the total number of repo-positive cells using the “spots” object and automatic detection parameters with local thresholding and background subtraction. Brain volume was determined by using the “surfaces” object on the Phalloidin channel to encompass the entire 3dimensional volume of the brain. Graphs of raw repo-positive counts per brain as well as glial density (repo-positive counts divided by brain volume) were created in Graphpad Prism (v9.4.1) software. Glial quantifications were performed using full Z-stacks of whole mount brains. Welch’s T-test was used for comparisons between control and *tau^R406W^* animals (n=9 animals per group). The significance threshold was set to p < 0.05. Error bars represent the 95% confidence interval.

#### Differential gene expression

Differential gene expression analyses were performed using Model-based Analysis of Single-cell Transcriptomics (MAST) for each cell cluster (Finak et al., 2015). In brief, generalized linear hurdle models were used to compute differential expression, where logistic regression was used to account for stochastic dropouts, and a gaussian linear model was fitted to predict gene expression levels. Differential expression was determined by a likelihood ratio test. We required that differentially expressed genes meet a significance threshold of BH-adjusted p-value < 0.05; absolute log2 fold-change > 0.1; and detectable (non-zero) expression in at least 10% of cells in the cluster. Cellular detection rate (CDR, fraction of genes reliably detected in each cell) was included as a covariate in all regression models, as in published protocols (Finak et al., 2015). CDR acts as a proxy for estimating the effect of dropout events, amplification efficiency, cell volume, and other extrinsic factors while performing expression-related regression analyses. Analyses of tau-induced differential expression also included age as a regression model covariate. Separately, aging-induced changes within each cell cluster were computed from control data (*elav-GAL4 / +*), comparing differential gene expression between day 1 and 10, and day 10 and 20. In order to evaluate robustness and replicability, cross-sectional, tau-induced differentially expressed genes were also computed in day 10 animals (tau vs control) and results were compared between the discovery and replication datasets. For differential expression of the human cell subclusters reported in Mathys et al. (2019), normalized counts between individuals with AD pathology (n=24) and low/no pathology (n=24) were compared using MAST for each cell subcluster as described above.

To assess cell type-specific differences in regulon gene expression levels (Figures 4C), we performed differential regulon analysis using base statistical packages in R. In brief, mean expression of gene members in each regulon were computed per cell, and cell type matched comparisons were made between control and *elav>tau^R406W^* using linear regression, including age as a covariate. Specifically, for each cell type, a likelihood ratio test compared the fit of a full model (Regulon Expression ~ Genotype + age) and a reduced model (Regulon Expression ~ age), evaluating the contribution of genotype to model fit. Significance was set at a BH-adjusted p-value < 0.05. Regulon log2-fold changes were computed for *elav>tau^R406W^* versus control mean expression in each cluster.

Overrepresentation analysis (ORA) of differentially expressed gene sets were completed using the R implementation of WEBGESTALT (Wang et al., 2013). The following databases were used: Gene Ontology (GO) biological processes, GO molecular functions, GO cellular component, KEGG, and Panther. Enrichment significance was defined by hypergeometric test, followed by p-value adjustment using the BH-procedure; significance was set at p < 0.05. ORA of the tau unique gene set (n=363 genes) was performed using gProfiler (Raudvere et al., 2019). The Gene Ontology (GO), Human phenotype ontology (HP), KEGG, miRTarBase (MIRNA), Transfac (TF), and WikiPathways (WP) databases were used for querying genes. The organism parameter was set to “dmelanogaster” and the “fdr” correction method was used to apply the BH multiple testing correction. A false discovery rate (FDR) < 0.05 was the threshold for significance.

#### Multiple regression with elastic net

To identify features driving cell vulnerability in our scRNAseq dataset, we pooled information across cell clusters by performing elastic net regression. For all clusters showing significant *elav>tau^R406W^* vs. control cell abundance changes, log2FC values were regressed on the cell type-specific mean expression for 183 regulons. Given our goal to identify factors that influence cell type-specific vulnerability, we focused on 8 cell clusters with significant cell loss (FDR < 0.05). In a secondary analysis, we repeated elastic net regression and considered a larger number of potential predictor variables including i) the 183 regulons (as above); ii) 2,793 unique GO, KEGG and Panther pathways found to be significantly enriched among *elav>tau^R406W^* differentially expressed genes; iii) 7 WGCNA modules altered in *elav>tau^R406W^* (Mangleburg et al., 2020); and iv) curated NFκB signaling pathways. In addition, we also considered a large number of v) cell cluster technical parameters as potential predictors, including normalized cell counts, mean tau transgene expression, sum of UMIs, mean percent mitochondrial reads, and number of tau-induced differentially expressed genes (age-adjusted). For this analysis, all computed variables (e.g. GO pathways, WGCNA modules, regulons, NFκB genes), were first averaged within each cell, then averaged across all cells in order to determine a mean value for each cell cluster. Cluster-level means for all gene sets were computed using pooled cell data from both *elav>tau^R406W^* and controls and all ages. For gene sets derived from ORA analyses, we restricted consideration to those differentially expressed genes driving enrichment. We generated a matrix consisting of rows for each cell cluster and columns with values / means for each potential predictor variable.

We used the *caret* and *glmnet* packages in R to organize the data and perform elastic net regularized regression. Alpha (ridge vs lasso characteristic) and lambda (shrinkage parameter) values were tuned in a 1,000 by 1,000 grid using repeated 3-fold cross validation in caret, and the average root mean squared errors (RMSE) from testing the partitions were used to assess model performance. 3-fold cross validation was repeated 100 times for all alpha-lambda pairs, using a different data fold split for each iteration in order to account for variability in model performance from random sample partitioning. The mean of all prediction errors was used to assess the final performance of each alpha-lambda pair, and we selected the model with the lowest root mean square error (RMSE). Lastly, we generated a ranklist of predictor variables for tau-induced cell abundance changes based on the magnitude of coefficients from the selected model (Figure S13B and Table S10).

#### Correlation with human snRNAseq datasets

Previously published human snRNAseq data (Mathys et al., 2019, and Grubman et al., 2019) were reprocessed and filtered using the identical pipeline, as describe above for *Drosophila* data. Raw gene counts were normalized using the NormalizeData function in Seurat v3. The resulting 70,634 filtered cells from the Mathys data were re-clustered using our pipeline above for visual representation in Figure 5; however, the cell cluster annotations from the original publication were preserved. 13,214 filtered cells from the Grubman data were similarly promoted for analysis. *Drosophila* orthologs of genes detected in human datasets were determined using the DRSC Integrated Ortholog Prediction Tool (DIOPT) (Hu et al., 2011), requiring a minimum DIOPT score threshold of 5 or greater. If more than 1 fly ortholog was identified, we selected the ortholog with either i) the highest DIOPT score, ii) the highest weighted DIOPT score, or iii) the highest ranked option (best score when mapped both forward and reverse). Thus, 5,630 or 4,145 human-fly gene ortholog pairs, respectively, were considered for cross-species analyses of the Mathys and Grubman datasets. We scaled normalized expression of each gene with mean = 0 and variance = 1. Cluster-level gene expression was computed by averaging scaled expression values from all cells. Subsequently, we performed Pearson correlation analysis for all cluster pairs to quantify transcriptional similarities between fly and human cell, examining pairwise correlation coefficients for all gene-orthologs across all clusters. For visualization, we generated heatmaps representing Pearson correlation coefficients by seriation with hierarchical clustering. Association statistics for human neuropathological traits (heatmap at top of Figure 5A) were repurposed directly from the published supplementary from Mathys et al. (2019). For quantification of overlap between human microglia and fly ensheathing glia, we examined conserved differentially expressed genes using the hypergeometric overlap test.

## SUPPLEMENTAL INFORMATION

Supplemental information includes 14 Supplemental Figures and 13 Supplemental Tables.

