## Supplemental Figures for "Tau polarizes an aging transcriptional signature to excitatory neurons and glia"

### Figure S1

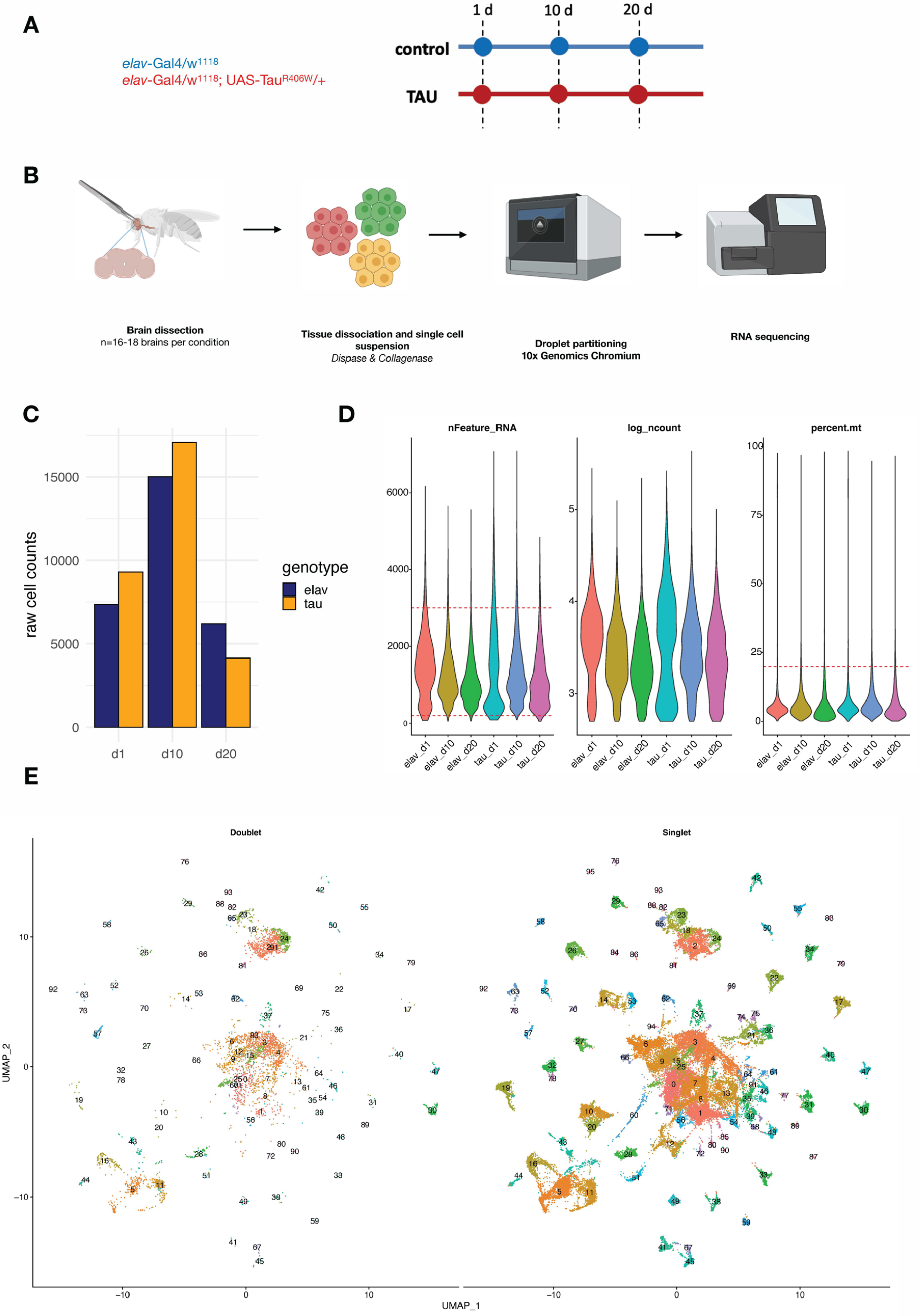
A
elav-Gal4/w1118
elav-Gal4/w1118; UAS-TauR406W/+
1 d
10 d
20 d
control
TAU
B
Brain dissection
n=16-18 brains per condition
Tissue dissociation and single cell suspension
Dispase & Collagenase
Droplet partitioning
10x Genomics Chromium
RNA sequencing
C
raw cell counts
genotype
elav
tau
d1
d10
d20
D
nFeature_RNA
log_ncount
percent.mt
elav_d1
elav_d10
elav_d20
tau_d1
tau_d10
tau_d20
E
Doublet
Singlet
UMAP_1
UMAP_2

**Figure S1. Study design and quality control metrics (Related to Figure 1).** (A) Schematic showing longitudinal study design for this study. Control (*elav-GAL4 / +*) and *elav>tau*$^{R406W}$ transgenic animals (*elav-GAL4 / +; UAS-tau*$^{R406W}$ */ +*) animals were aged to 3 timepoints: 1, 10, and 20 days. (B) For each library, 16-18 brains were dissected from the cuticle and pooled for dissociation into a single cell suspension. Cells were partitioned into single cell droplets using the 10x Genomics Chromium platform for library preparation, and completed libraries were sequenced using the Illumina NovaSeq 6000. Figures were generated using BioRender. (C) Plot shows the number of cells captured in each library for *elav-GAL4* controls (elav) and *elav>tau*$^{R406W}$ (tau) at each timepoint: day 1 (d1), day 10 (d10), or day 20 (d20). (D) Violin plots display scRNAseq library quality control metrics, including the number of unique genes captured (nFeature_RNA), log(number of UMIs) (log_ncount), and % mitochondrial reads (percent.mt). Cell filtering cutoffs are denoted by red dashed lines. (E) Unsupervised clustering of cells after filtering by DoubletFinder, showing cells classified as doublets or multiplets (left) and singlets (right). A total of 48,111 cells classified as singlets were used for downstream analyses.

### Figure S2

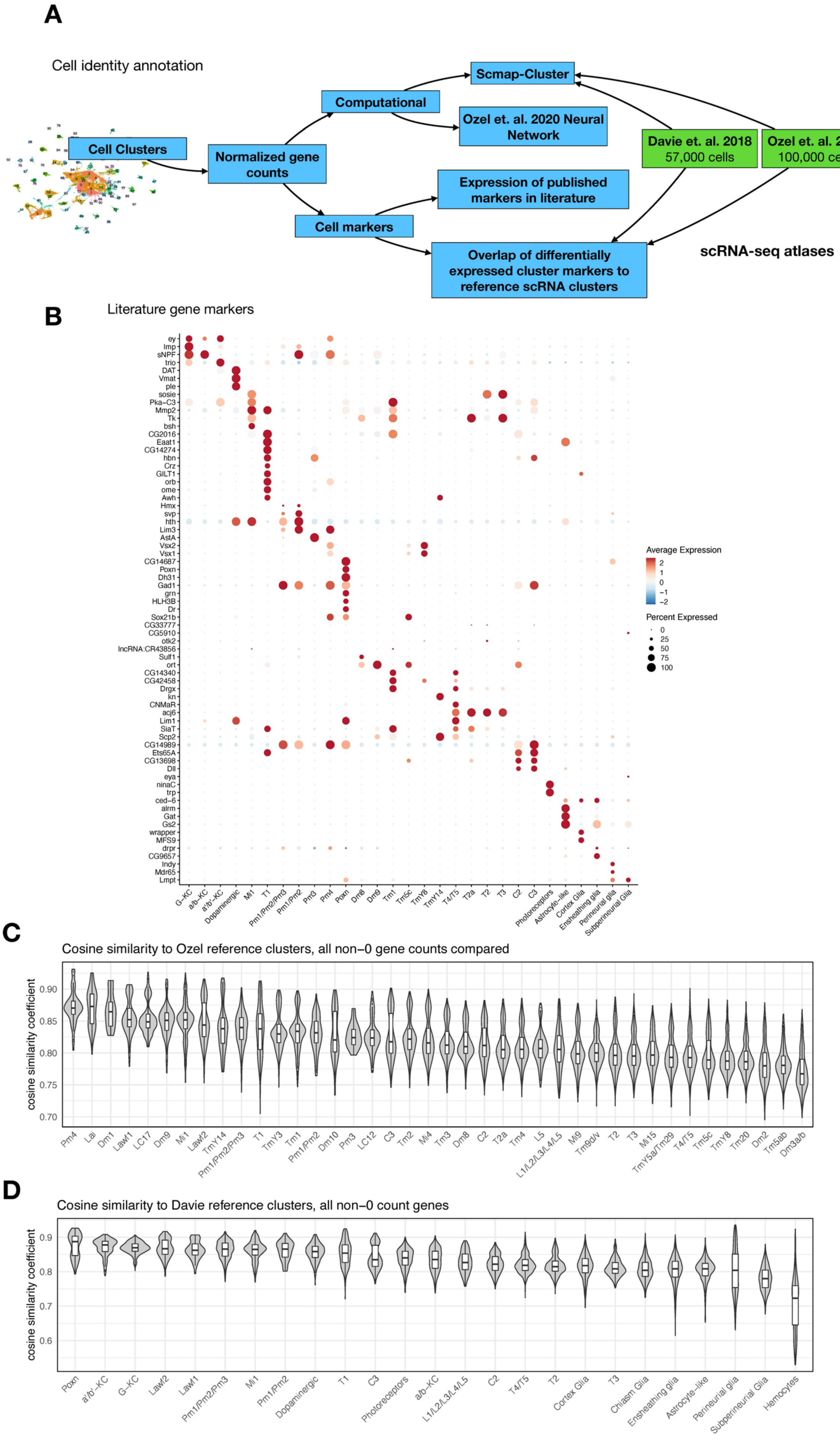
A
Cell identity annotation
Cell Clusters
Normalized gene counts
Computational
Scmap-Cluster
Ozel et. al. 2020 Neural Network
Cell markers
Expression of published markers in literature
Overlap of differentially expressed cluster markers to reference scRNA clusters
Davie et. al. 2018 57,000 cells
Ozel et. al. 2020 100,000 cells
scRNA-seq atlases
B
Literature gene markers
Average Expression
Percent Expressed
C
Cosine similarity to Ozel reference clusters, all non−0 gene counts compared
cosine similarity coefficient
D
Cosine similarity to Davie reference clusters, all non−0 count genes
cosine similarity coefficient

**Figure S2. Annotating cell identities for 96 cell clusters across 48,111 cells from Drosophila brains (Related to Figure 1).** (A) Schematic shows the cell identity annotation pipeline utilized for this study. We leveraged both published scRNA-seq atlases from Davie et al (2018) and Ozel et al (2020) as well as other well-established cell type markers. Our strategy included (i) correlation-based approach (scmap) using the 2 brain atlases as reference, (ii) a 2-layer neural network classifier from Ozel et al (2020) to identify optic lobe neurons, and (iii) differential expression of cell-specific marker genes. (B) Expression of cell-specific markers across annotated cell identities. Normalized expression for each gene is scaled (Z-transformed) across all cell clusters and represented in a blue-red color scale. Percent of cells in each cluster that have detectable (non-zero) gene expression is represented by dot size. (C) Correlation analysis (cosine similarity) of shared, non-dropout, genes from individual annotated cells in our dataset to cluster-level means of the corresponding cell identity in Ozel et al (2020). Violin plot representing distribution of cosine similarity scores of annotated cells in our dataset to their (available) corresponding reference cluster in Ozel et al (2020). The median similarity coefficient for cell-to-reference pairings ranged from 0.87 to 0.76. (D) Identical correlation analysis as described in (C), applied with reference clusters from Davie et al (2018). The median similarity coefficient for cell-to-reference pairings ranged from 0.9 to 0.72.

### Figure S3

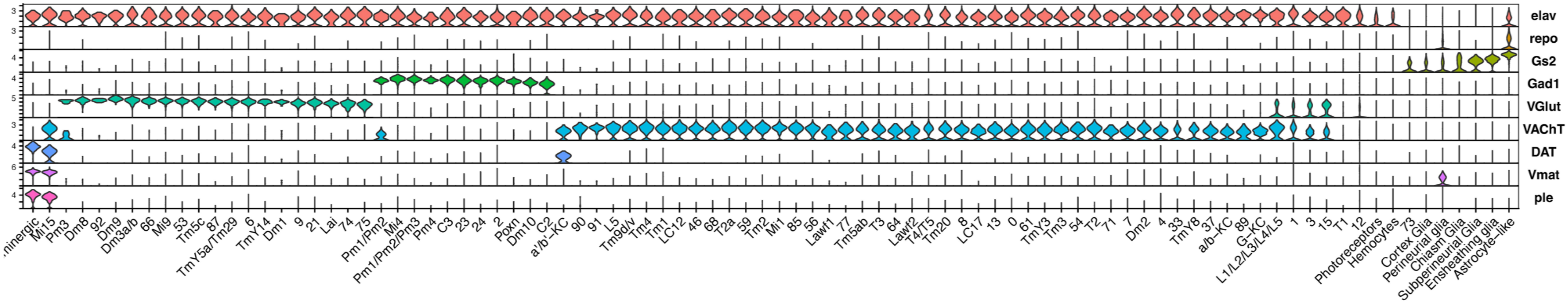

elav
repo
Gs2
Gad1
VGlut
VAChT
DAT
Vmat
ple
Mi15
Pm3
Dm8
92
Dm9
Dm3a/b
66
Mi9
53
Tm5c
87
TmY5a/Tm29
6
TmY14
Dm1
9
21
Lai
74
75
Pm1/Pm2
Mi4
Pm1/Pm2/Pm3
Pm4
C3
23
24
2
Poxn
Dm10
C2
a'/b'-KC
90
91
L5
Tm9d/v
Tm4
Tm1
LC12
46
68
T2a
59
Tm2
Mi1
85
56
Lawf1
77
Tm5ab
T3
64
Lawf2
T4/T5
Tm20
8
LC17
13
0
61
TmY3
Tm3
54
T2
71
7
Dm2
4
33
TmY8
37
a/b-KC
89
G-KC
L1/L2/L3/L4/L5
1
3
15
T1
12
Photoreceptors
Hemocytes
73
Cortex Glia
Perineurial glia
Chiasm Glia
Subperineurial Glia
Ensheathing glia
Astrocyte-like

**Figure S3. Normalized gene expression of general cell type markers across all defined cell clusters (Related to Figure 1).** Violin plot of general cell type markers for all cell clusters, as described in Figure 1B.

### Figure S4

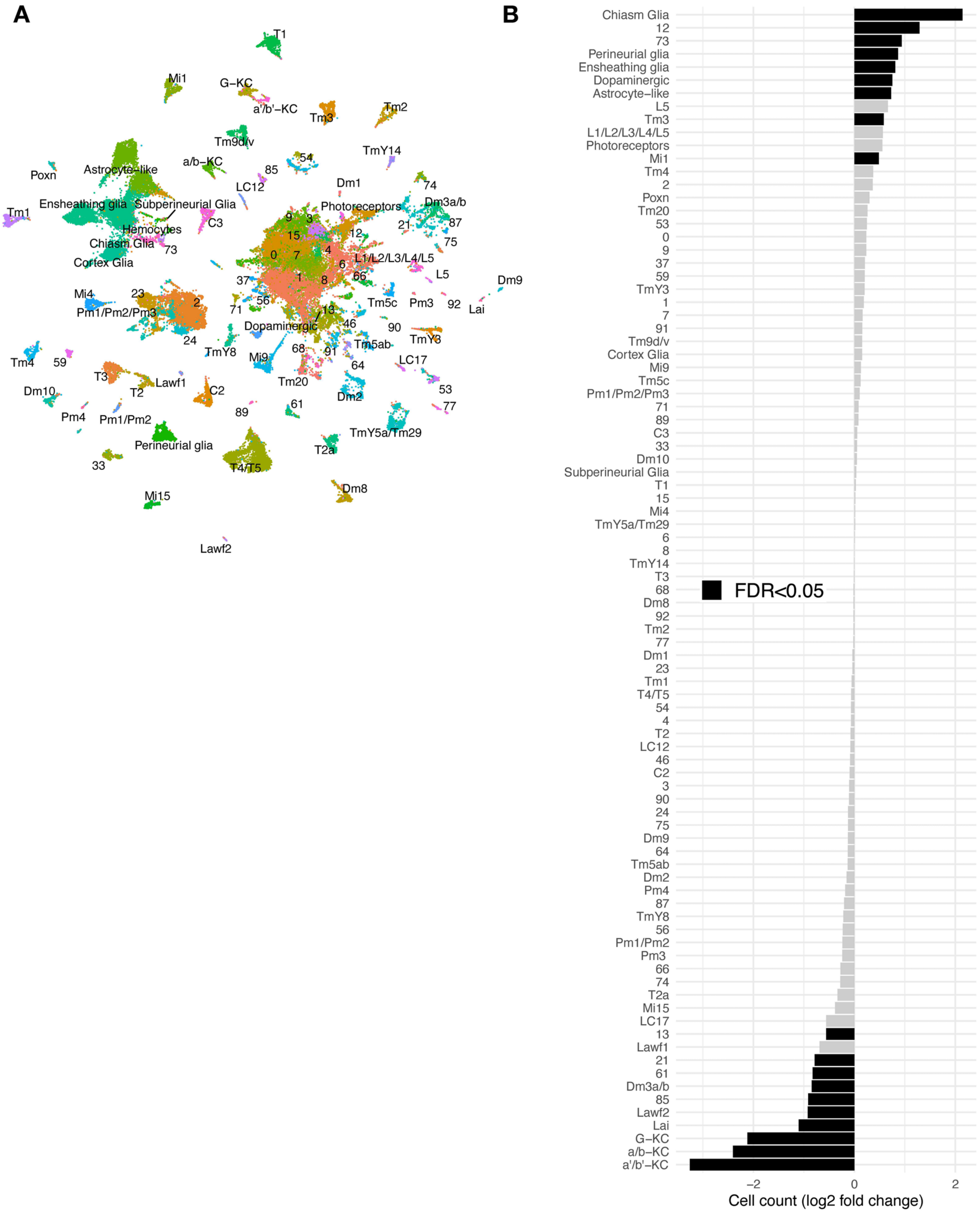
A
B
T1
Mi1
G–KC
a'/b'–KC
Tm3
Tm2
Tm9d/v
a/b–KC
54
TmY14
85
Poxn
Astrocyte–like
LC12
Dm1
74
Ensheathing glia
Subperineurial Glia
Photoreceptors
Dm3a/b
Tm1
C3
9
3
21
87
Hemocytes
15
12
Chiasm Glia
73
4
75
0
7
6
L1/L2/L3/L4/L5
Cortex Glia
1
8
66
L5
37
Dm9
Mi4
23
2
56
Tm5c
Pm3
92
Lai
Pm1/Pm2/Pm3
71
13
Dopaminergic
46
90
TmY3
24
TmY8
Mi9
68
91
Tm5ab
LC17
Tm4
59
64
T3
Lawf1
Tm20
53
Dm10
T2
C2
Dm2
77
61
89
Pm4
Pm1/Pm2
TmY5a/Tm29
Perineurial glia
T2a
33
T4/T5
Dm8
Mi15
Lawf2
Chiasm Glia
12
73
Perineurial glia
Ensheathing glia
Dopaminergic
Astrocyte–like
L5
Tm3
L1/L2/L3/L4/L5
Photoreceptors
Mi1
Tm4
2
Poxn
Tm20
53
0
9
37
59
TmY3
1
7
91
Tm9d/v
Cortex Glia
Mi9
Tm5c
Pm1/Pm2/Pm3
71
89
C3
33
Dm10
Subperineurial Glia
T1
15
Mi4
TmY5a/Tm29
6
8
TmY14
T3
68
Dm8
92
Tm2
77
Dm1
23
Tm1
T4/T5
54
4
T2
LC12
46
C2
3
90
24
75
Dm9
64
Tm5ab
Dm2
Pm4
87
TmY8
56
Pm1/Pm2
Pm3
66
74
T2a
Mi15
LC17
13
Lawf1
21
61
Dm3a/b
85
Lawf2
Lai
G–KC
a/b–KC
a'/b'–KC
FDR<0.05
−2
0
2
Cell count (log2 fold change)

**Figure S4. Additional scRNAseq from 3 *tau*$^{R406W}$ and 3 control libraries at day 10 post-eclosion (Related to Figures 2 and 3).** (A) UMAP plot showing the 69,128 cells comprising the scRNAseq replication dataset from 10-day-old control (*elav-GAL4 / +*) and *elav>tau*$^{R406W}$ (*elav-GAL4 / +; UAS-tau*$^{R406W}$ */ +*) flies. Cell cluster names are consistent with that used in the discovery dataset (Figure 1). (B) Plots shows tau-triggered cell abundance changes, based on $\log_2$-fold change of normalized cell counts in the replication dataset. 19 clusters have statistically significant cell abundance changes [False Discovery Rate (FDR) < 0.05; black bars]. Of note, in the replication dataset, hemocytes were present in only very low numbers, and therefore were not included in cell abundance analyses.

### Figure S5

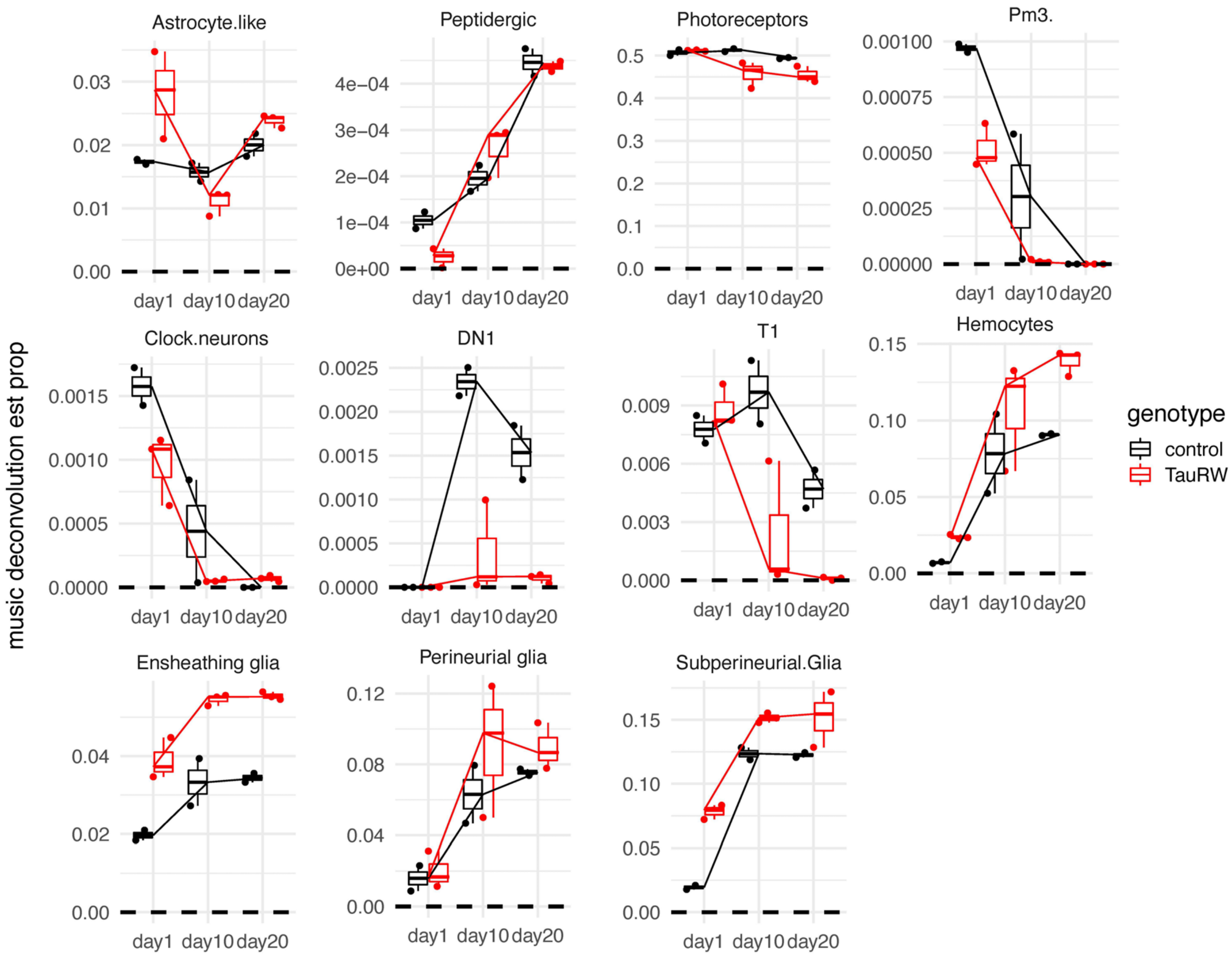

Astrocyte.like
Peptidergic
Photoreceptors
Pm3.
Clock.neurons
DN1
T1
Hemocytes
Ensheathing glia
Perineurial glia
Subperineurial.Glia
music deconvolution est prop
day1
day10
day20
genotype
control
TauRW

**Figure S5. Estimation of cell proportions by deconvolution of bulk-tissue RNA-sequencing (Related to Figure 2).** Cell proportions (y-axis) are shown for selected cell types of interest, based on analysis of bulk-tissue RNAseq from control (black, n=2) (*elav-GAL4 / +*) and *elav>tau*$^{R406W}$ (red, n=3) (*elav-GAL4 / +; UAS-tau*$^{R406W}$ */ +*) flies (Mangleburg et al., 2020). A line is drawn through the median of each condition. There is an elevation in the estimated proportions of astrocyte-like, ensheathing, and perineurial glia, recapitulating observations in the scRNAseq cell abundance analysis.

### Figure S6

**A**

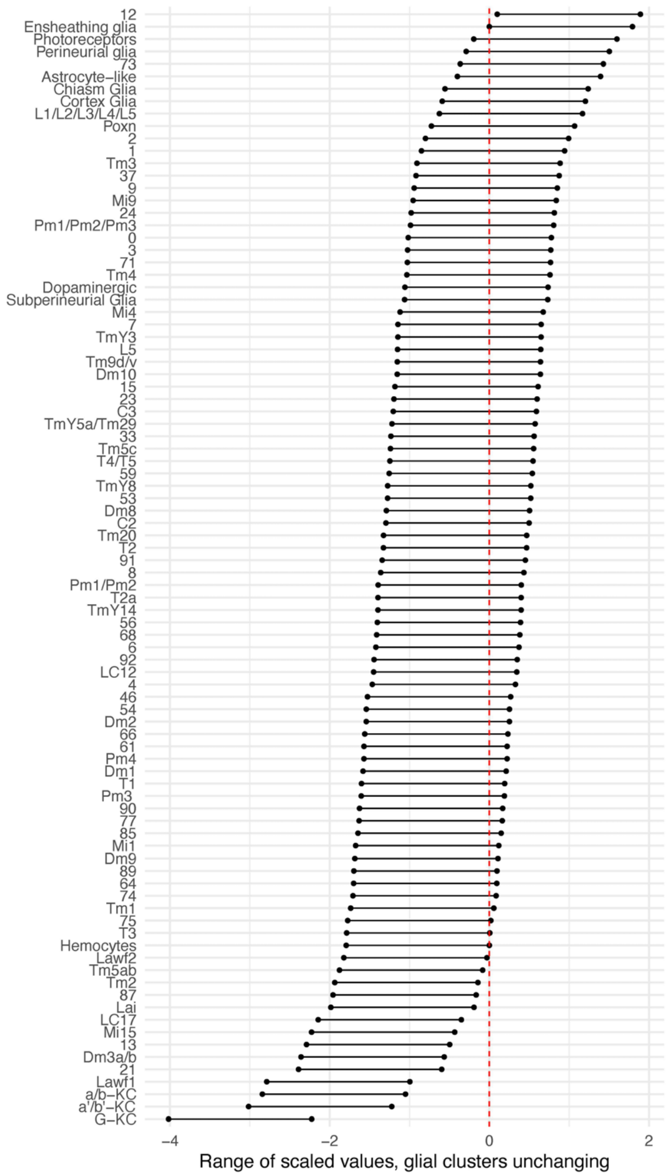
12
Ensheathing glia
Photoreceptors
Perineurial glia
73
Astrocyte–like
Chiasm Glia
Cortex Glia
L1/L2/L3/L4/L5
Poxn
2
1
Tm3
37
9
Mi9
24
Pm1/Pm2/Pm3
0
3
71
Tm4
Dopaminergic
Subperineurial Glia
Mi4
7
TmY3
L5
Tm9d/v
Dm10
15
23
C3
TmY5a/Tm29
33
Tm5c
T4/T5
59
TmY8
53
Dm8
C2
Tm20
T2
91
8
Pm1/Pm2
T2a
TmY14
56
68
6
92
LC12
4
46
54
Dm2
66
61
Pm4
Dm1
T1
Pm3
90
77
85
Mi1
Dm9
89
64
74
Tm1
75
T3
Hemocytes
Lawf2
Tm5ab
Tm2
87
Lai
LC17
Mi15
13
Dm3a/b
21
Lawf1
a/b–KC
a'/b'–KC
G–KC
−4
−2
0
2
Range of scaled values, glial clusters unchanging

**B**

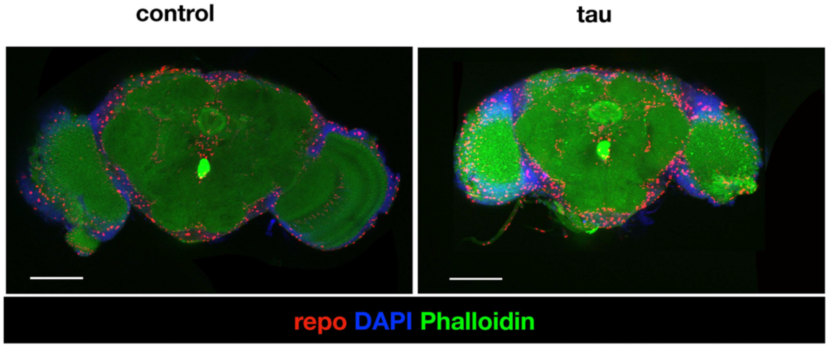
control
tau
repo DAPI Phalloidin

**Figure S6. Adjusted tau-triggered cell abundance changes (Related to Figure 2).** (A) In order to adjust for proportional changes, the log2 fold-change value for 7 cell clusters (Ensheathing glia, Perineurial glia, Astrocyte-like glia, Cortex glia, Chiasm glia, Subperineurial glia, and Hemocytes) was iteratively subtracted from the cell abundance estimates for all other clusters, establishing a confidence interval. Following adjustment, 14 decreasing and 1 increasing cell types are highlighted. All other cell type clusters have fold-change estimates overlapping zero. (B) Whole-mount immunofluorescence of adult brains from 10-day-old flies, including control (*elav-GAL4 / +*) and *elav>tau*$^{R406W}$ (*elav-GAL4 / +; UAS-tau*$^{R406W}$ */ +*) flies. Composite of 10 confocal sections is shown, from co-staining for glia (Anti-Repo, red) along with nuclei (DAPI, blue) and actin (phalloidin, green). Scale bar = 100 microns.

### Figure S7

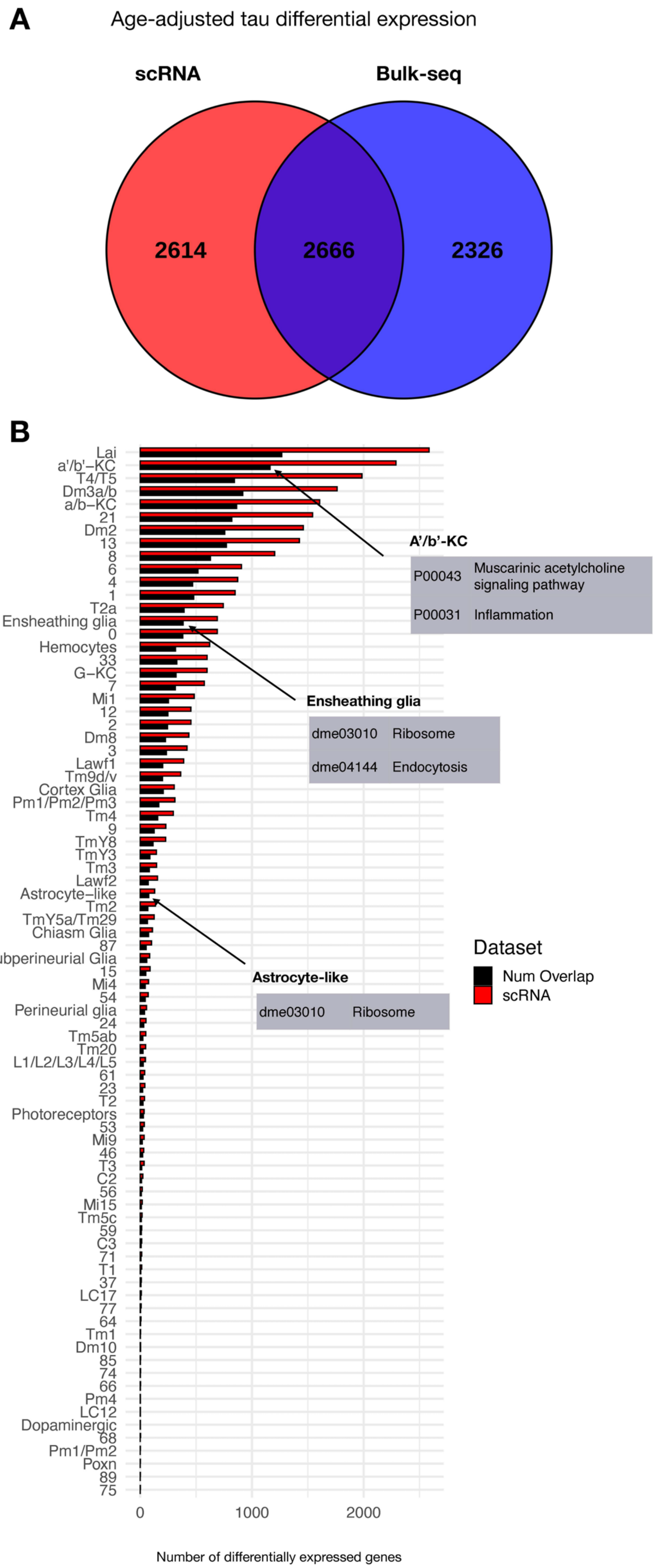
A
Age-adjusted tau differential expression
scRNA
Bulk-seq
2614
2666
2326
B
A'/b'-KC
P00043 Muscarinic acetylcholine signaling pathway
P00031 Inflammation
Ensheathing glia
dme03010 Ribosome
dme04144 Endocytosis
Astrocyte-like
dme03010 Ribosome
Dataset
Num Overlap
scRNA
Number of differentially expressed genes
0
1000
2000
Lai
a'/b'−KC
T4/T5
Dm3a/b
a/b−KC
21
Dm2
13
8
6
4
1
T2a
Ensheathing glia
0
Hemocytes
33
G−KC
7
Mi1
12
2
Dm8
3
Lawf1
Tm9d/v
Cortex Glia
Pm1/Pm2/Pm3
Tm4
9
TmY8
TmY3
Tm3
Lawf2
Astrocyte−like
Tm2
TmY5a/Tm29
Chiasm Glia
87
Subperineurial Glia
15
Mi4
54
Perineurial glia
24
Tm5ab
Tm20
L1/L2/L3/L4/L5
61
23
T2
Photoreceptors
53
Mi9
46
T3
C2
56
Mi15
Tm5c
59
C3
71
T1
37
LC17
77
64
Tm1
Dm10
85
74
66
Pm4
LC12
Dopaminergic
68
Pm1/Pm2
Poxn
89
75

**Figure S7. Overlap between tau-induced adult brain gene expression changes between *Drosophila* scRNAseq and bulk-tissue RNA-sequencing (Related to Figure 3).** (A) Venn diagram illustrates the number of tau-induced differentially expressed genes in bulk (blue) vs. single-cell (red) RNAseq. These complementary analyses consider identical genotypes and timepoints, including control (*elav-GAL4 / +*) and *elav>tau*$^{R406W}$ (*elav-GAL4 / +; UAS-tau*$^{R406W}$ */ +*) transgenic flies profiled at 1-, 10-, and 20-days. Both regression analyses similarly adjust for age. (B) Plot shows the number of tau-induced differentially expressed genes per cell cluster from scRNAseq data (red), and the number of overlapping, differentially expressed genes from bulk tissue RNAseq (black). Overall, genes that are uniquely differentially expressed in the scRNAseq data (n=2,614 genes) are restricted to fewer cell clusters, whereas shared gene expression changes (n=2,666 genes) are expressed more broadly that are observed in both platforms (shared differentially expressed genes, 2,666 genes). Further, genes that are uniquely differentially expressed in bulk RNAseq data, where sequencing reads are not diluted across individual cells, tend to be expressed at lower levels when compared to those that are shared.

### Figure S8

A

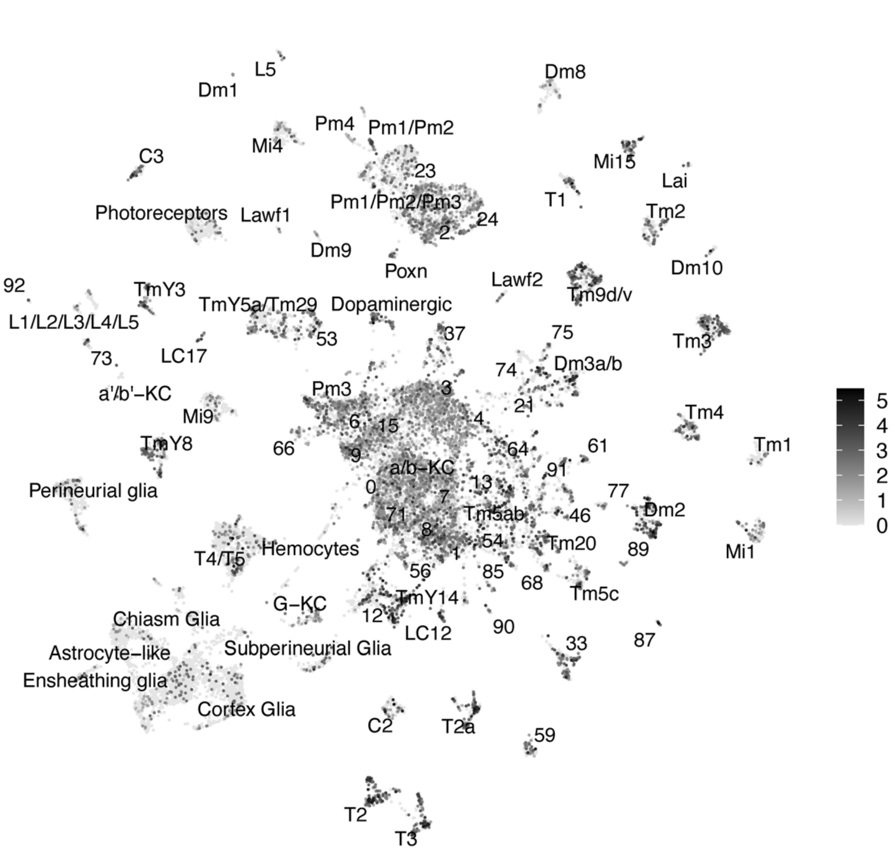
mapt
Dm8
L5
Dm1
Pm4
Pm1/Pm2
Mi4
Mi15
C3
Lai
23
Pm1/Pm2/Pm3
T1
Photoreceptors
Lawf1
Tm2
24
2
Dm9
Dm10
Poxn
Lawf2
92
TmY3
Tm9d/v
TmY5a/Tm29
Dopaminergic
L1/L2/L3/L4/L5
75
53
37
Tm3
73
LC17
Dm3a/b
74
a'/b'−KC
Pm3
3
21
Mi9
Tm4
4
15
6
TmY8
Tm1
66
64
61
9
91
a/b−KC
0
Perineurial glia
13
77
7
71
Tm5ab
46
Dm2
8
54
Tm20
89
Mi1
T4/T5
Hemocytes
1
56
85
68
Tm5c
G−KC
TmY14
Chiasm Glia
12
LC12
90
87
33
Astrocyte−like
Subperineurial Glia
Ensheathing glia
Cortex Glia
C2
T2a
59
T2
T3
5
4
3
2
1
0

B

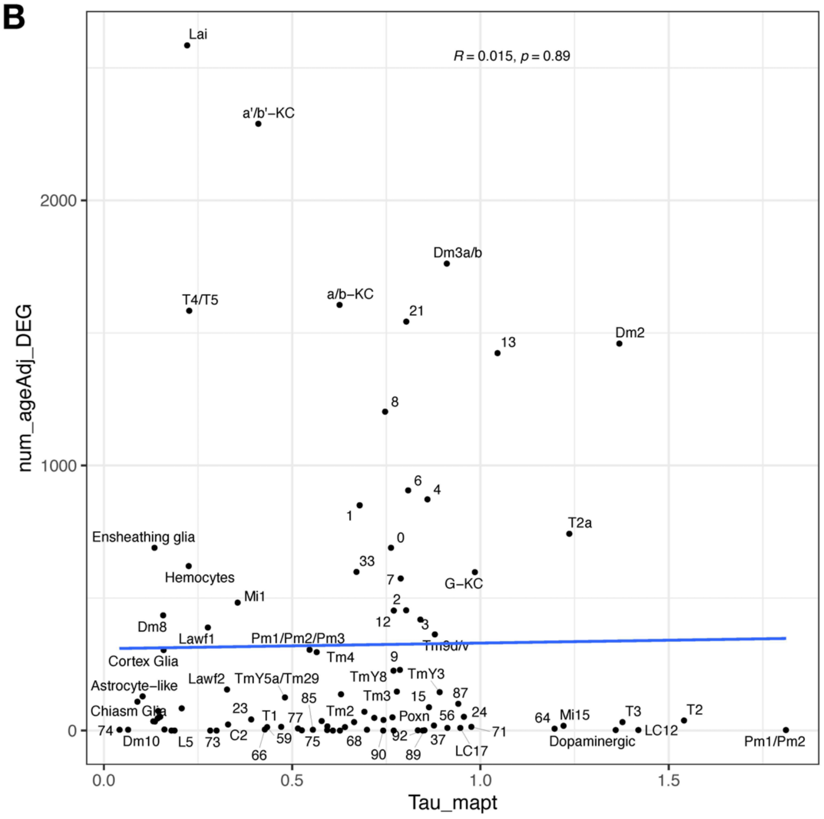
$R = 0.015, p = 0.89$
num_ageAdj_DEG
Tau_mapt
2000
1000
0
0.0
0.5
1.0
1.5
Lai
a'/b'−KC
Dm3a/b
T4/T5
a/b−KC
21
13
Dm2
8
6
4
1
T2a
Ensheathing glia
0
33
Hemocytes
7
G−KC
Mi1
2
Dm8
12
3
Lawf1
Pm1/Pm2/Pm3
Tm9d/v
Cortex Glia
Tm4
9
TmY8
TmY3
Astrocyte−like
Lawf2
TmY5a/Tm29
Tm3
15
87
85
Chiasm Glia
23
T1
77
Tm2
Poxn
56
24
64
Mi15
T3
T2
74
Dm10
L5
73
C2
66
59
75
68
90
92
89
37
LC17
71
Dopaminergic
LC12
Pm1/Pm2

**Figure S8. Expression of the *MAPT* transgene (Related to Figure 3).** (A) Uniform manifold approximation and projection (UMAP) plot showing widespread *MAPT* transgene expression across in *elav>tau*$^{R406W}$ (*elav-GAL4 / +; UAS-tau*$^{R406W}$ */ +*) animals. The pan-neuronal *elav-GAL4* driver induces widespread expression of tau. (B) Plot showing overall poor Pearson correlation (R=0.015, p=0.89) between number of tau-induced gene expression changes (y-axis) and the mean *MAPT* expression level per cell cluster (x-axis).

### Figure S9

A

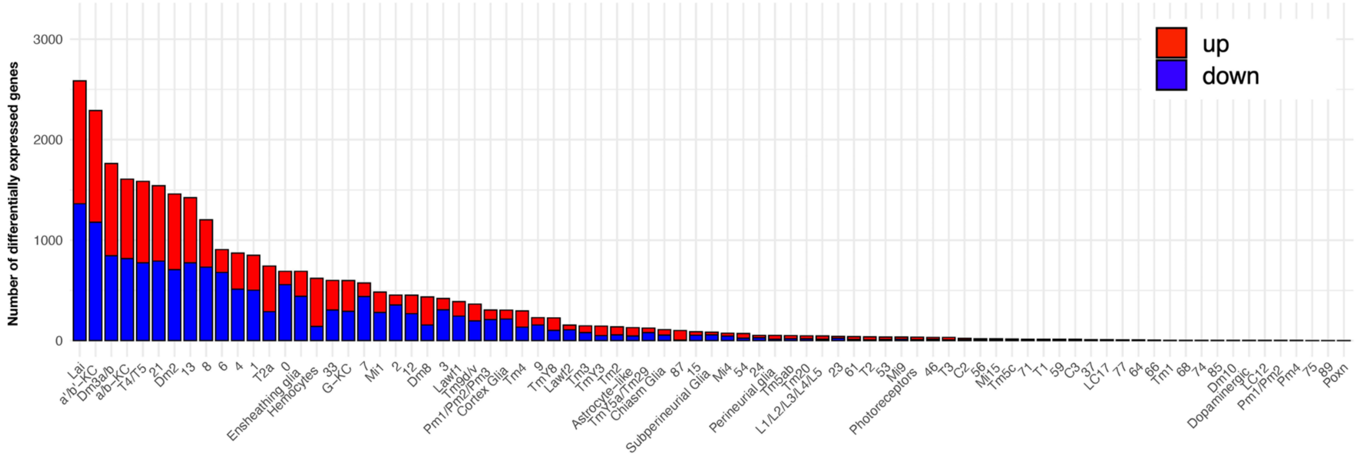

Number of differentially expressed genes
3000
2000
1000
0
up
down


B

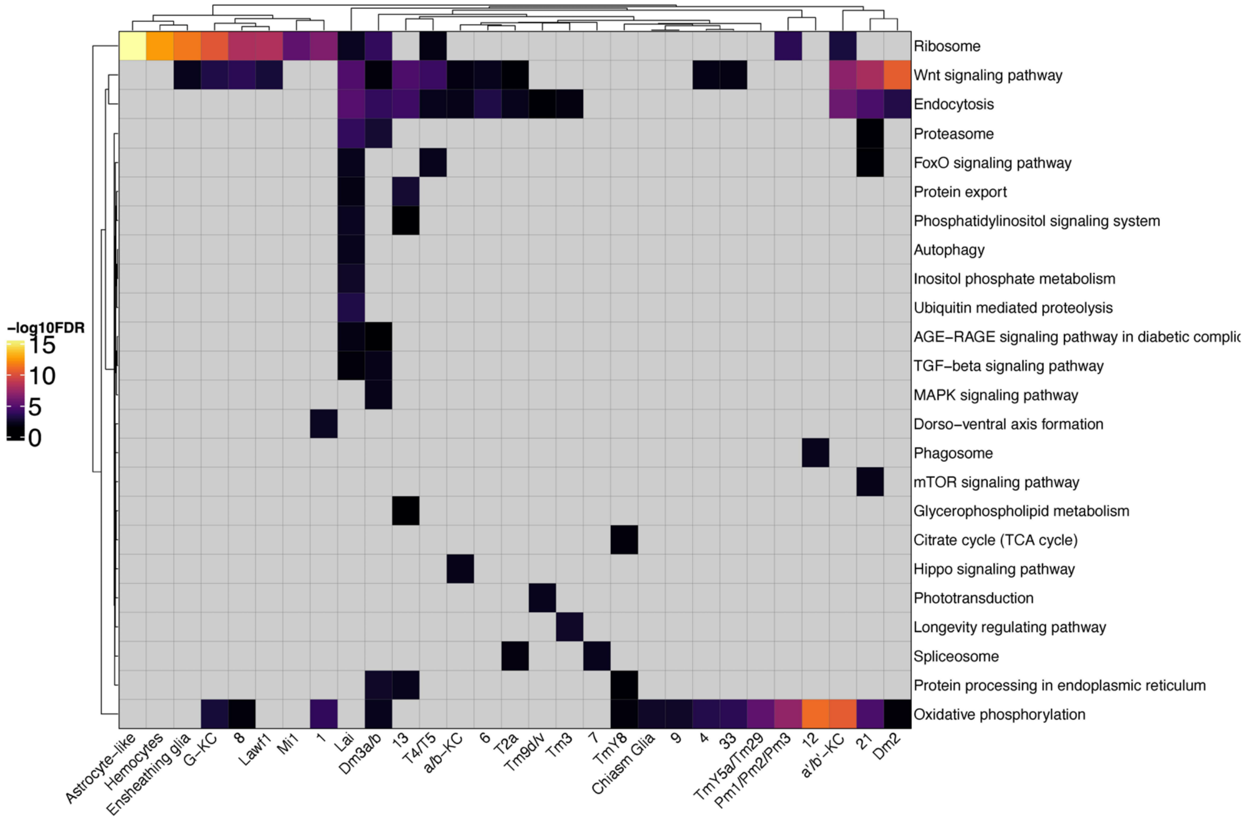

-log10FDR
15
10
5
0
Ribosome
Wnt signaling pathway
Endocytosis
Proteasome
FoxO signaling pathway
Protein export
Phosphatidylinositol signaling system
Autophagy
Inositol phosphate metabolism
Ubiquitin mediated proteolysis
AGE-RAGE signaling pathway in diabetic complic
TGF-beta signaling pathway
MAPK signaling pathway
Dorso-ventral axis formation
Phagosome
mTOR signaling pathway
Glycerophospholipid metabolism
Citrate cycle (TCA cycle)
Hippo signaling pathway
Phototransduction
Longevity regulating pathway
Spliceosome
Protein processing in endoplasmic reticulum
Oxidative phosphorylation
Astrocyte-like
Hemocytes
Ensheathing glia
G-KC
8
Lawf1
Mi1
1
Lai
Dm3a/b
13
T4/T5
a/b-KC
6
T2a
Tm9d/v
Tm3
7
TmY8
Chiasm Glia
9
4
33
TmY5a/Tm29
Pm1/Pm2/Pm3
12
a'/b'-KC
21
Dm2

**Figure S9. Tau-induced differential gene expression analysis and functional enrichment (Related to Figure 3).** (A) The number of tau-induced, differentially expressed genes are shown following adjustment for aging, but highlighting up- (red) versus down- (blue) regulated genes. Data presented is otherwise same as that shown in Figure 3B, based on comparisons of control (*elav-GAL4 / +*) and *elav>tau*$^{R406W}$ (*elav-GAL4 / +; UAS-tau*$^{R406W}$ */ +*). (B) Heatmap shows significant KEGG terms from functional enrichment analysis of tau-induced differentially expressed genes, including pathways that are actively in cell-type specific vs. more global patterns. Non-significant test results are shown in gray, whereas positive results [hypergeometric test, False Discovery Rate (FDR) $< 0.05$] are shaded based on significance level [-log10(FDR)].

### Figure S10

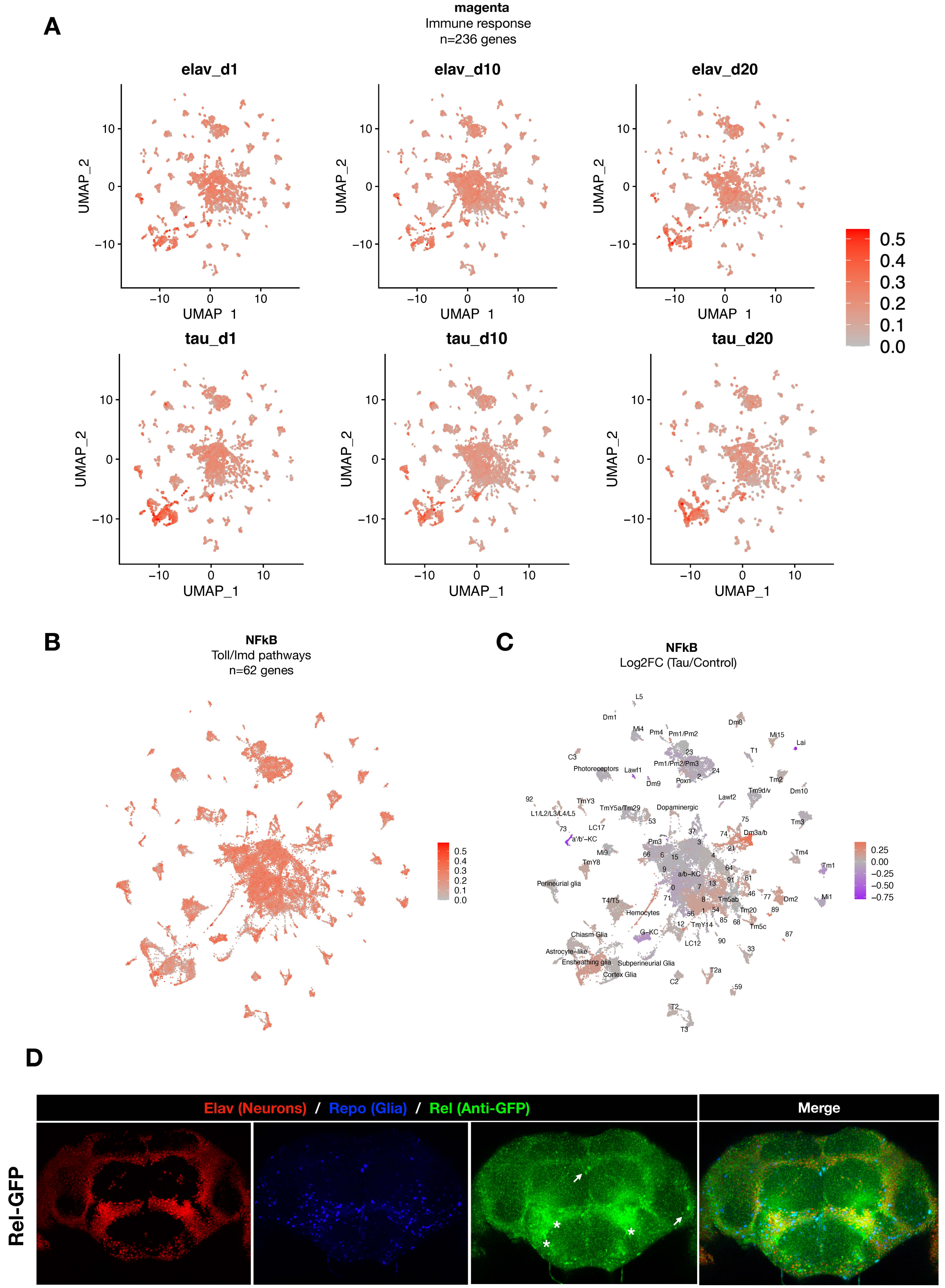

A
magenta
Immune response
n=236 genes
elav_d1
elav_d10
elav_d20
tau_d1
tau_d10
tau_d20
UMAP_1
UMAP_2
B
NFkB
Toll/Imd pathways
n=62 genes
C
NFkB
Log2FC (Tau/Control)
D
Elav (Neurons) / Repo (Glia) / Rel (Anti-GFP)
Merge
Rel-GFP

**Figure S10. Expression of immune response and NFkB genes in *Drosophila* brain (Related to Figure 4).** (A) Plots showing mean expression of the immune response gene coexpression module (n=236 genes), based on analyses of scRNAseq data stratified by genotype (control vs. *elav>tau*$^{R406W}$) or age (1-, 10-, or 20-days). Innate immune signaling appears to be broadly expressed across brain cell types for all conditions. (B) Plot shows mean overall normalized expression by cell cluster among n=62 curated NFkB signaling pathway genes (see Methods for full list). In this plot, gene expression was averaged across both *elav>tau*$^{R406W}$ and control cells. Results are similar to that seen for the immune response coexpression model (Figure 4A). (C) Plot shows $log_2$ fold-change mean expression per cell cluster for the same 62-gene NFkB signaling mediators, based on comparisons between *elav>tau*$^{R406W}$ and control flies. Results are similar to that seen for the immune response coexpression model (Figure 4B). (D) Experimental confirmation of Relish expression in adult *Drosophila* brains. Whole mount immunofluorescence of adult brains from Rel-GFP flies, in which the endogenous Relish protein harbors an amino-terminal GFP tag in homozygozity in an otherwise wildtype genetic background (*y, w; PBac{GFP.FPTB-Rel}VK00037*). Rel-GFP (Anti-GFP, green) were stained for neuronal nuclei (anti-Elav, red), glia (anti-Repo, blue). The asterisks and arrows denote Rel expression/localization to neuronal and glial nuclei, respectively.

### Figure S11

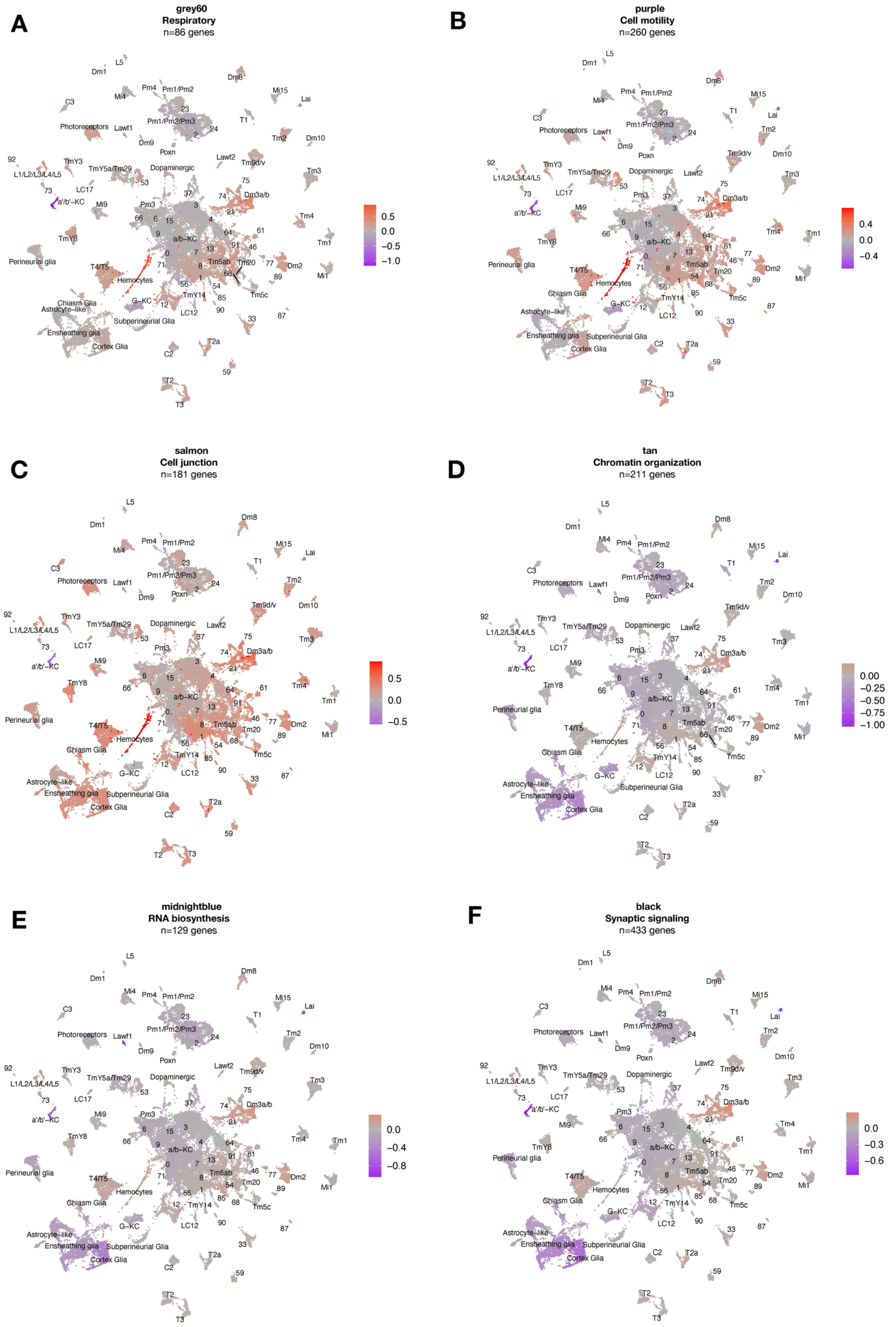
A
grey60
Respiratory
n=86 genes
0.5
0.0
−0.5
−1.0
B
purple
Cell motility
n=260 genes
0.4
0.0
−0.4
C
salmon
Cell junction
n=181 genes
0.5
0.0
−0.5
D
tan
Chromatin organization
n=211 genes
0.00
−0.25
−0.50
−0.75
−1.00
E
midnightblue
RNA biosynthesis
n=129 genes
0.0
−0.4
−0.8
F
black
Synaptic signaling
n=433 genes
0.0
−0.3
−0.6

**Figure S11. Cell-type specific expression of tau-induced gene coexpression modules (Related to Figure 4).** Uniform manifold approximation and projection (UMAP) plots show gene expression changes for several published, tau-triggered gene coexpression modules. Color scale shows mean gene expression changes per cell cluster (log2FC), based on scRNAseq data from *elav>tau*$^{R406W}$ (*elav-GAL4 / +; UAS-tau*$^{R406W}$ */ +*) and control (*elav-GAL4 / +*) flies. Modules are functionally enriched for genes involved in (A) respiration, (B) cell motility, (C) cell junctions, (D) chromatin organization, (E) RNA biosynthesis, and (F) synaptic signaling, as previously described (Mangleburg et al, 2020).

### Figure S12

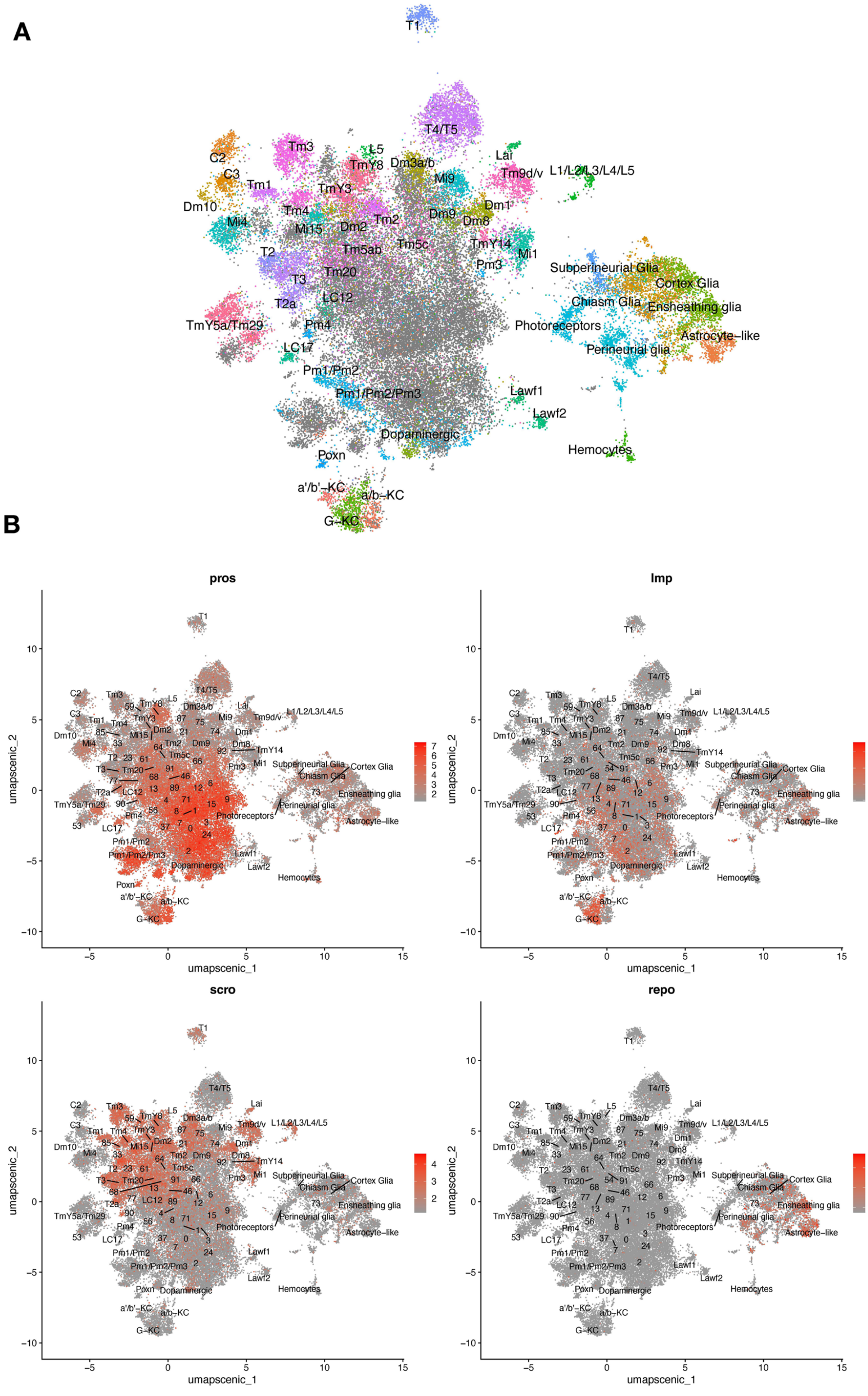
A
T1
T4/T5
C2
Tm3
L5
TmY8
Dm3a/b
Lai
Tm9d/v
L1/L2/L3/L4/L5
C3
Tm1
Mi9
TmY3
Dm10
Tm4
Dm1
Mi4
Mi15
Dm2
Tm2
Dm9
Dm8
TmY14
Tm5c
Mi1
T2
Tm5ab
Pm3
Subperineurial Glia
Cortex Glia
T3
Tm20
T2a
LC12
Chiasm Glia
Ensheathing glia
TmY5a/Tm29
Pm4
Photoreceptors
Astrocyte−like
LC17
Perineurial glia
Pm1/Pm2
Pm1/Pm2/Pm3
Lawf1
Lawf2
Dopaminergic
Hemocytes
Poxn
a'/b'−KC
a/b−KC
G−KC
B
pros
Imp
scro
repo
umapscenic_1
umapscenic_2

**Figure S12. Unsupervised clustering based on regulon coexpression networks (Related to Figures 1 and 4).** (A) Uniform manifold approximation and projection (UMAP) plots show relationships among 48,111 cells based on 183 regulons (cell-level regulon activity scores). (B) Cell-specific expression of brain cell marker genes. Whereas *Pros* and *Imp* expression appears widespread, *scro* show more restricted expression to optic lobe neurons, and *repo* expression is limited to glia.

### Figure S13

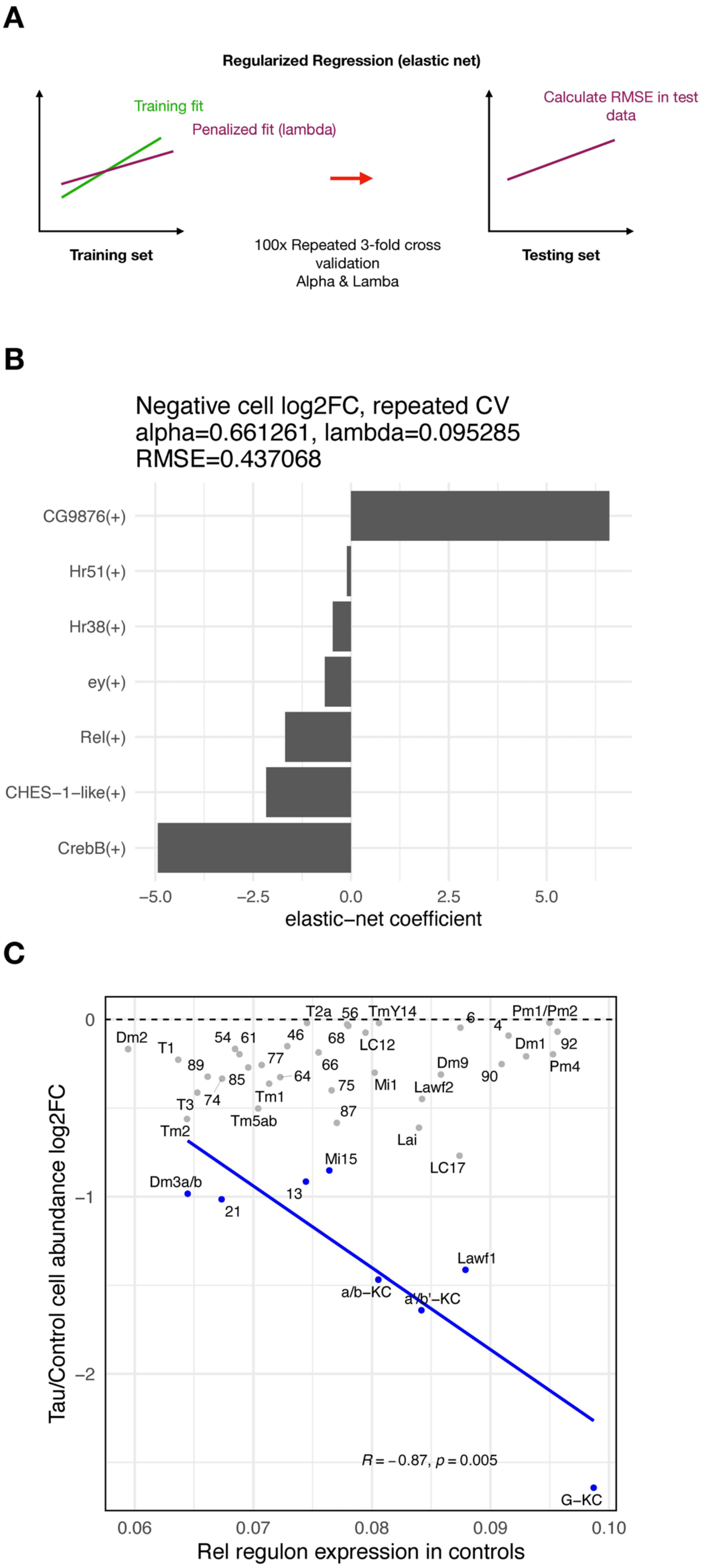

A
Regularized Regression (elastic net)
Training fit
Penalized fit (lambda)
Training set
100x Repeated 3-fold cross validation
Alpha & Lamba
Calculate RMSE in test data
Testing set
B
Negative cell log2FC, repeated CV
alpha=0.661261, lambda=0.095285
RMSE=0.437068
CG9876(+)
Hr51(+)
Hr38(+)
ey(+)
Rel(+)
CHES−1−like(+)
CrebB(+)
−5.0
−2.5
0.0
2.5
5.0
elastic−net coefficient
C
Tau/Control cell abundance log2FC
Rel regulon expression in controls
0
−1
−2
0.06
0.07
0.08
0.09
0.10
T2a
56
TmY14
6
4
Pm1/Pm2
Dm2
T1
54
61
46
68
LC12
Dm1
92
89
77
66
Dm9
Pm4
85
64
75
Mi1
90
Lawf2
T3
74
Tm1
87
Tm5ab
Tm2
Lai
Mi15
LC17
Dm3a/b
13
21
Lawf1
a/b−KC
a'/b'−KC
R = −0.87, p = 0.005
G−KC

**Figure S13. Regulons associated with tau-induced cell vulnerability (Related to Figure 4).** (A) Schematic showing analytic strategy to identify regulon expression networks that predict tau-triggered cell loss. We implemented elastic net regression to examine the relation between regulon expression (predictor variable) and cell abundance changes (response variable) for clusters showing significant tau-induced cell abundance changes. 3-fold cross validation was repeated 100 times for hyperparameter tuning (alpha-lambda). (B) Plot showing the regression coefficients for prioritized regulons (out of 183 total) that predict cell abundance changes in *elav>tau*$^{R406W}$ flies. The Rel regulon was the 3$^{rd}$ ranked predictor for the severity of neuronal loss. (C) We replotted Figure 4D, showing the relation between Rel regulon expression, *but restricted to control animals*, and tau-induced cell abundance changes. Among clusters with significant, tau-induced cell loss (denoted in blue, FDR<0.05), cell abundance change remained inversely correlated with Rel regulon expression (Pearson correlation: R = -0.87, p=0.005).

### Figure S14

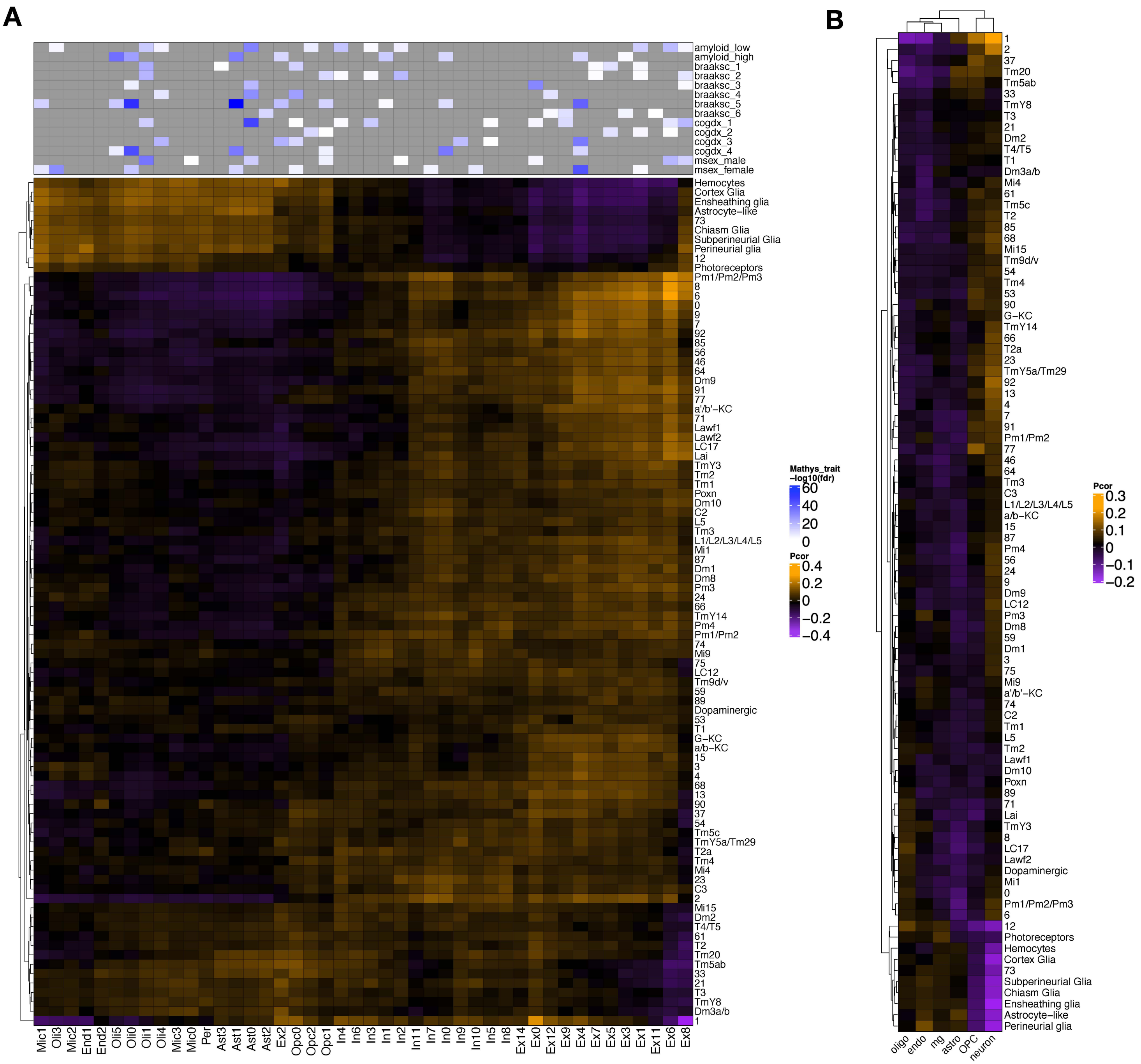
A
B
amyloid_low
amyloid_high
braaksc_1
braaksc_2
braaksc_3
braaksc_4
braaksc_5
braaksc_6
cogdx_1
cogdx_2
cogdx_3
cogdx_4
msex_male
msex_female
Mathys_trait
-log10(fdr)
60
40
20
0
Pcor
0.4
0.2
0
−0.2
−0.4
Hemocytes
Cortex Glia
Ensheathing glia
Astrocyte−like
73
Chiasm Glia
Subperineurial Glia
Perineurial glia
12
Photoreceptors
Pm1/Pm2/Pm3
8
6
0
9
7
92
85
56
46
64
Dm9
91
77
a'/b'−KC
71
Lawf1
Lawf2
LC17
Lai
TmY3
Tm2
Tm1
Poxn
Dm10
C2
L5
Tm3
L1/L2/L3/L4/L5
Mi1
87
Dm1
Dm8
Pm3
24
66
TmY14
Pm4
Pm1/Pm2
74
Mi9
75
LC12
Tm9d/v
59
89
Dopaminergic
53
T1
G−KC
a/b−KC
15
3
4
68
13
90
37
54
Tm5c
TmY5a/Tm29
T2a
Tm4
Mi4
23
C3
2
Mi15
Dm2
T4/T5
61
T2
Tm20
Tm5ab
33
21
T3
TmY8
Dm3a/b
1
Mic1
Oli3
Mic2
End1
End2
Oli5
Oli0
Oli1
Oli4
Mic3
Mic0
Per
Ast3
Ast1
Ast0
Ast2
Ex2
Opc0
Opc2
Opc1
In4
In6
In3
In1
In2
In11
In7
In0
In9
In10
In5
In8
Ex14
Ex0
Ex12
Ex9
Ex4
Ex7
Ex5
Ex3
Ex1
Ex11
Ex6
Ex8
Pcor
0.3
0.2
0.1
0
−0.1
−0.2
oligo
endo
mg
astro
OPC
neuron

**Figure S14. Cross-species gene expression correlation of all *Drosophila* cell clusters in this study (Related to Figure 5).** (A) Heatmap shows Pearson correlation of gene expression (5,700 conserved, orthologous genes in total) between annotated cell clusters from *Drosophila* (rows) and human postmortem brain (columns; Mathys et al., 2019). Compared with Figure 5A, this plot includes the non-annotated *Drosophila* cell clusters. (B) Similar heatmap was constructed based on 4,145 conserved orthologs from *Drosophila* and an independent human AD case-control snRNAseq dataset (Grubman et al., 2019). Annotated human cell types include oligodendrocytes (oligo), endothelial cells (endo), microglia (mg), astrocytes (astro), oligodendrocyte precursor cells (OPC), and neurons.
